# Drosophila ß_Heavy_-Spectrin is required in polarized ensheathing glia that form a diffusion-barrier around the neuropil

**DOI:** 10.1101/2021.07.21.453164

**Authors:** Nicole Pogodalla, Holger Kranenburg, Simone Rey, Silke Rodrigues, Albert Cardona, Christian Klämbt

## Abstract

In the central nervous system (CNS), functional tasks are often allocated to distinct compartments. This is also evident in the insect CNS where synapses and dendrites are clustered in distinct neuropil regions. The neuropil is separated from neuronal cell bodies by ensheathing glia, which as we show using dye injection experiments forms an internal diffusion barrier. We find that ensheathing glial cells are polarized with a basolateral plasma membrane rich in phosphatidylinositol-(3,4,5)-triphosphate (PIP_3_) and the Na^+^/K^+^-ATPase Nervana2 (Nrv2) that abuts an extracellular matrix formed at neuropil-cortex interface. The apical plasma membrane is facing the neuropil and is rich in phosphatidylinositol-(4,5)-bisphosphate (PIP_2_) that is supported by a sub-membranous ß_Heavy_-Spectrin cytoskeleton. *ß_Heavy_-spectrin* mutant larvae affect ensheathing glial cell polarity with delocalized PIP_2_ and Nrv2 and exhibit an abnormal locomotion which is similarly shown by ensheathing glia ablated larvae. Thus, polarized glia compartmentalizes the brain and is essential for proper nervous system function.

## Introduction

The complexity of the nervous system is not only defined by intricate neuronal networks but is also reflected by the cellular diversity of the different glial cells found throughout the nervous system. The number of functions attributed to the different glial cells is steadily growing and ranges from guiding and instructing functions during early brain development to synapse formation and plasticity, and general ion and metabolite homeostasis (Yildirim et al., 2018; Zuchero and Barres, 2015).

One of the key functions of glial cells lies in the establishment of boundaries that help compartmentalizing neuronal computing. A very tight boundary is the blood-brain barrier. In all invertebrates as well as in primitive vertebrates this barrier is established by glial cells, whereas higher vertebrates transferred this function to endothelial cells (Bundgaard and Abbott, 2008; Carlson et al., 2000; Schirmeier and Klämbt, 2015). Smaller signaling compartments are established by astrocytes that tile the synaptic areas in the brain (Freeman, 2010; Magistretti and Allaman, 2018; Tsacopoulos and Magistretti, 1996). Oligodendrocytes insulate single segments of up to 20 different axons by forming myelin sheets around them (Simons and Nave, 2016). Moreover, oligodendrocyte progenitor cells act as innate immune cells and participate in scar formation separating intact brain tissue from the injured site (Fernandez-Castaneda and Gaultier, 2016).

Invertebrates have a less complex nervous system and glial cells are clearly outnumbered by neurons. In the ant *Harpegnathos saltator* 27 % of all brain cells are of glial nature, whereas in the fruit fly *Drosophila melanogaster* just 10 % of all neural cells are of glial nature (Davie et al., 2018; Sheng et al., 2020). As in primitive vertebrates, glial cells form the blood-brain barrier and organize the entire metabolite traffic into and out of the nervous system and also integrate external stimuli such as circadian rhythms with general brain functions (Hindle et al., 2017; Limmer et al., 2014; Volkenhoff et al., 2015; Zhang et al., 2018) All neuronal cell bodies are situated beneath the blood-brain barrier and are surrounded by cortex glial cells. These cells regulate neuroblast division, nurture neurons, and phagocytose dying neurons (Cabirol-Pol et al., 2017; Coutinho-Budd et al., 2017; Etchegaray et al., 2016; Hilu-Dadia et al., 2018; Kis et al., 2015; Liu et al., 2017; 2015; Nakano et al., 2019).

Fly neurons project their dendrites and axons into the central neuropil. This morphologically distinct structure is separated from all neuronal cell bodies by the neuropil associated glia, which comprises astrocyte-like glial cells and ensheathing glial cells (Ito et al., 1995; Otto et al., 2018; Peco et al., 2016; Stacey et al., 2010; Stork et al., 2014). Astrocyte-like cells form numerous fine and highly branched processes that infiltrate the neuropil. As in vertebrates they tile the synaptic neuropil into discrete units. They express a number of neurotransmitter transporters to clear synaptic spillover although they do not form frequent intimate contacts with individual synapses and in addition are able to secrete gliotransmitters (Ma et al., 2016; MacNamee et al., 2016; Stork et al., 2014).

The neuropil encasing ensheathing glia comprises two distinct classes that can be classified by morphological as well as molecular criteria (Davie et al., 2018; Kremer et al., 2017; Otto et al., 2018; Peco et al., 2016). In the larval nervous system, just four ensheathing glial cells are formed in each hemineuromer. Two of them embrace the neuropil and do not wrap around individual axons. In contrast, the other two larval ensheathing glial cells found in each abdominal hemineuromere exhibit a more complex morphology. In part they encase the neuropil as the other two ensheathing glial cells but in addition they also enwrap axons between the CNS/PNS boundary and the neuropil and are thus called ensheathing/wrapping glial cells (Otto et al., 2018; Peco et al., 2016). In the larva, the organization of the neuropil is relatively simple, but the adult CNS shows complex regional compartmentalization (Hartenstein et al., 2008; Zheng et al., 2018). This is reflected by an increase in the number of ensheathing glial cells that are generated during pupal stages and as in larval stages fall into two morphological classes (Kremer et al., 2017; Omoto et al., 2015).

Several studies have already shed some light on the functional roles of ensheathing glia in the fly nervous system. First, it was demonstrated that ensheathing glial cells remove neuronal debris after injury utilizing the Draper pathway (Doherty et al., 2009; Hilu-Dadia et al., 2018; Lu et al., 2017). In addition to this immune and surveillance function, ensheathing glia can participate in neuronal signaling. Mutant analysis indicates the sulfite oxidase Shopper needs to be expressed by ensheathing glia to regulate glutamate homeostasis in the neuropil (Otto et al., 2018). Likewise, the Excitatory amino acid transporter 2 (Eaat2) functions in ensheathing glia to modulate sleep in the adult (Stahl et al., 2018). In addition, cell type specific knockdown experiments demonstrate that the voltage-gated potassium channel encoded by the gene *seizure* (*sei*) is required in ensheathing glia to protect flies from acute heat-induced seizures (Hill et al., 2019).

Given the apparent role of ensheathing glia in forming neuronal compartments and modulating their function, we initiated a comprehensive analysis of this - as we found - highly polarized cell type. The plasma membrane of the ensheathing glia facing the neuropil is rich in the phospholipid PIP_2_ and is supported by sub-membranous ß_Heavy_-Spectrin. Thus, it can be assumed that the apical cell domain is oriented towards the neuropil. The basolateral plasma membrane exhibits an accumulation of the Na^+^ / K^+^ ATPase Nervana 2 (Nrv2) suggesting that sodium ion dependent transport is polarized in these cells. In addition, basally localized Integrins anchor the ensheathing glia to a specific extracellular matrix (ECM) formed at the interface of neuropil and CNS cortex. To study the functional relevance of ensheathing glia, we ablated these cells using a newly established split Gal4 driver. The absence of ensheathing glia triggers a compensatory growth of astrocyte-like cells which however, does not completely restore an internal diffusion barrier normally established by the ensheathing glia. Animals lacking ensheathing glia show abnormal larval locomotion and reduced longevity. Reduction of ß_Heavy_-Spectrin expression affects ensheathing glia cell morphology leads to mislocalization of PIP_2_. Likewise, ß_Heavy_-Spectrin mutants show a locomotor phenotype as observed following ensheathing glia ablation. In conclusion, we show that separation of the neuropil by polarized ensheathing glia is required for nervous system function.

## Results

### Ensheathing glia encase the neuropil throughout development

Four ensheathing glial cells are formed in each hemineuromere during mid embryogenesis and are associated with the neuropil (Peco et al., 2016). Two of these cells only encase the neuropil, whereas the two other also wrap axons connecting the peripheral nerves with the neuropil (ensheathing/wrapping glial cells, **Figure 1**) (Beckervordersandforth et al., 2008; Kremer et al., 2017; Peco et al., 2016). To target the larval ensheathing glial cells, we compared the activity of the previously used Gal4 lines (*NP6520-Gal4*, *56F03-Gal4*, *nrv2-Gal4* and *83E12-Gal4*) and found the *83E12*-Gal4 driver as the most specific one for both larval and adult CNS (Kremer et al., 2017; Li et al., 2014; Otto et al., 2018; Peco et al., 2016). Cell counts and 3D reconstructions (see below) suggest that *83E12-Gal4* is active in all larval ensheathing glial cells. Since no 3D reconstruction has been conducted for the adult TEM volume we cannot exclude that some *83E12-Gal4* negative ensheathing glial cells exist.

**Figure 1.**
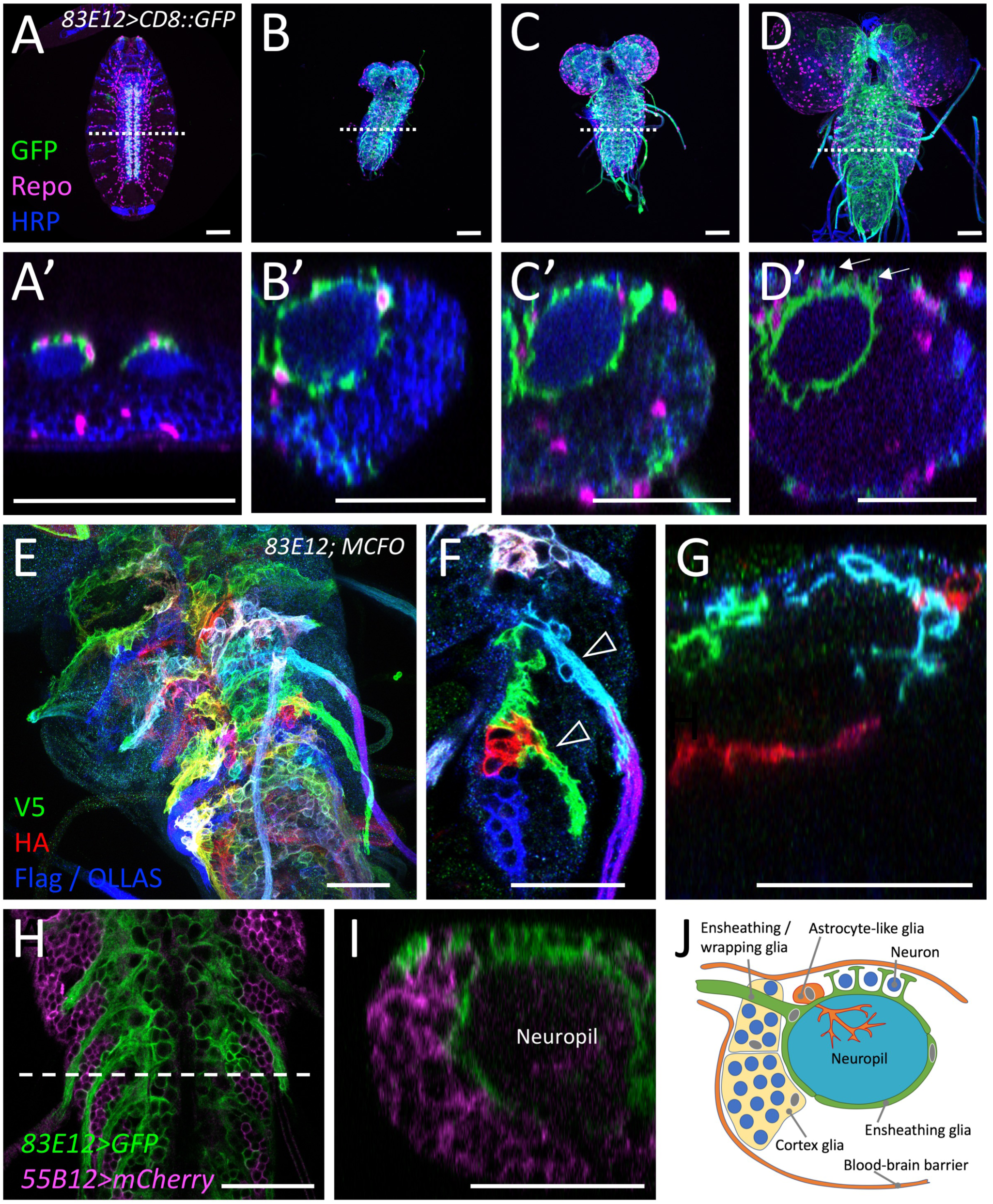
Development of ensheathing glia (see also supplementary Figure 1) (A-D) Dissected larval CNS of increasing age with the genotype [*83E12-Gal4, UAS-CD8::GFP*], stained for GFP (green), Repo (magenta) and neuronal membranes (anti-HRP, blue). The positions of the orthogonal section shown in (A’-D’) is indicated by a dashed line. Scale bars are 50 µm, anterior is up. (A,A’) In a stage 16 embryo ensheathing glial cells have not yet covered the neuropil (arrowheads). (B,B) First instar larval CNS. (C,C’) Second instar larval CNS. (D,D’) CNS of a wandering third instar larva. The arrows point towards dorsal protrusions of the ensheathing glia engulfing dorsal neurons. (E) MCFO labeling of ensheathing glia in third instar larva stained for the expression of V5 (green), HA (red), and FLAG and OLLAS epitopes (blue). *flp* expression was induced for one hour during first instar larval stage. Note that ensheathing glia tile the ventral nerve cord. (F) Two distinct wrapping/ensheathing glia cells cover the nerve root and part of the neuropil (arrowheads). (G) Ensheathing glia occupy specific territories in the neuropil. (H,I) Third instar larval nerve cord with the genotype [*55B12-Gal4, 83E12-LexA, UAS-CD8::mCherry, LexAop-GFP*]. All cortex glia cells are labelled by mCherry expression (magenta). Ensheathing glial cells are labelled by GFP expression (green). The dashed line indicates the position of the orthogonal view shown in (I). (J) Schematic view on a cross section through a hemineuromere indicating the position of the different glial cells. Astrocyte-like cells and ensheathing glial cells localize close to the neuropil and the axons connecting the neuropil with the periphery. The cortex glia covers lateral and ventral neuronal cell bodies.

During embryonic stages, the ensheathing glia initially cover the dorsal domain of the neuropil (**Figure 1A,A’**). In first instar larvae, thin processes of the ensheathing glia begin to encase the neuropil (**Figure 1B,B**’). Notably, small gaps between individual ensheathing glial processes are still detectable in the second instar larva (**Figure 1C,C’**). In the third larval instar, ensheathing glia appear to form a closed case around the neuropil. The ensheathing glia also send processes around the dorsally located neuronal cell bodies (**Figure 1D**’, **Figure S1**). Multi-color flipout (MCFO) labelling demonstrates that ensheathing glia processes tile the neuropil (**Figure 1E-G,J**) to separate it from the CNS cortex. The cortex glia can be visualized using the Gal4 drivers *55B12-Gal4; NP2222 and NP0577* (Awasaki et al., 2008; Otto et al., 2018). These drivers showed that no cortex glial cells reside at the dorsal surface of the ventral nerve cord (**Figure 1H-J**). This notion is further corroborated by split-GFP experiments where GFP is reconstituted at the baso-lateral boundary between cortex and neuropil but not at the dorsal surface of the ventral nerve cord (**Figure S1A-F’**). Here, several large glutamatergic neurons are found that appear to be encased by processes of the ensheathing glia **(Figure S1G-L**). In conclusion, the ensheathing glia can wrap axons as they connect with peripheral organs, associate with dorsal neurons, and encase the neuropil **(Figure 1J)**.

### Electron microscopic analysis of larval ensheathing glia

To further study the morphological characteristics of the ensheathing glia, we analyzed serial section transmission electron microscopy (ssTEM) data sets of a first instar larval ventral nerve cord (L1) (Ohyama et al., 2015). We annotated all neuropil associated glial cells in the abdominal neuromeres A1-A8/9 using CATMAID (Saalfeld et al., 2009) (**Figure 2**). In total, 159 neuropil-associated glial cells were identified that based on their typical morphology could be assigned to one of the three classes of neuropil-associated glial cells (Peco et al., 2016): astrocyte-like glial cells, ensheathing glia and ensheathing/wrapping glia (**Figure 2A,B**). The morphological characteristics of the different glial subtypes are already evident in first larval instar and become more pronounced in third instar larval brain (**Figure 2C,D**). Astrocyte-like glial cells were identified by a prominent process extending into the neuropil proximal to the astrocyte nucleus (MacNamee et al., 2016; Stork et al., 2014). In the abdominal ventral nerve cord of the L1 volume, 96 astrocyte-like glial cells were identified matching the previously known numbers (Peco et al., 2016; Stork et al., 2014).

28 ensheathing glial cells were identified in the abdominal neuromeres 1-7 (**Figure 2A**). These glial cells extend thin membrane sheaths along the face of the neuropil (**Figure 2E**). The nuclei of the central ensheathing glial cell in each neuromere is usually associated with the dorso-ventral channel (Ito et al., 1995). The other ensheathing glia is found at a ventral position at the neuropil cortex interface (see also(Peco et al., 2016)).

**Figure 2.**
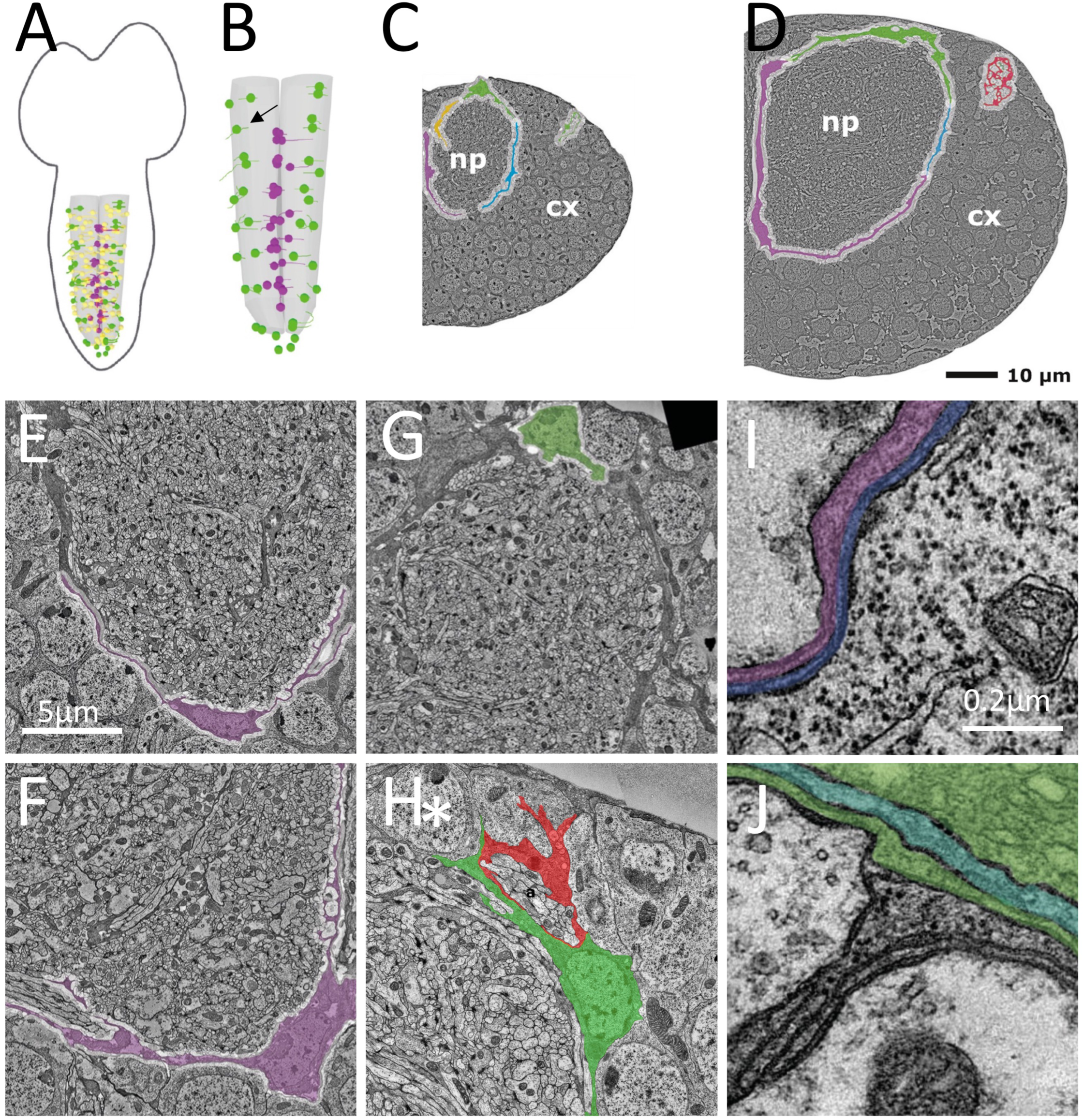
Larval development of the ensheathing glia. (A,B) Schematic view on a first instar larval central nervous system. All neuropil associated glial cells were annotated in a serial section TEM volume. Yellow dots indicate the position of the nuclei of astrocyte-like cells, magenta dots indicate the positions of the ensheathing glial cell nuclei, green dots indicate the positions of the ensheathing / wrapping glial nuclei. The arrow points to a segmental position where only one instead of two ensheathing wrapping glial cell was identified. (C) Exemplary section through the abdominal 4 showing the position of three different ensheathing glial cells around the neuropil. It was not determined whether the red marked glial processes belong to the blue or green ensheathing wrapping glial cell. (D) Section through the fourth abdominal neuromere of a third instar larval ventral nerve cord. Note that ensheathing glia completely cover the neuropil. Scale bar (C,D): 10 µm, (E-H): 5µm, (I,J): 0.2 µm. (E) Ensheathing glia in a first and (F) in a third instar larva. (G) Ensheathing / wrapping glial cells in a first and (H) in a third instar larva. Note the thin processes that engulf cell bodies of dorsally located neurons (asterisk). (I) On the ventral face of the neuropil an ensheathing glial cell and a cortex glial cell form a two-layered sheath between cortex and neuropil. (J) A highly multilayered glial cell layer is found at the dorsal face of the neuropil.

33 cells were annotated as ensheathing/wrapping glial cells, which encase the neuropil and wrap axons in the nerve root. However, this is not very pronounced in the first instar stage, yet (**Figure 2G**). With one exception (**Figure 2B**), two cells, both located at the dorsal aspect of the neuropil, are associated with each intersegmental nerve root in neuromeres 1-7 (**Figure 2B**). The fused A8/A9 nerve has two nerve roots associated with two ensheathing/wrapping glial cells, each (**Figure 2B**).

To determine whether the overall morphology of ensheathing glial cells changes during development, we annotated the ensheathing glia in three abdominal neuromeres (A1-A3) of a third instar larval brain (L3) (Valdes-Aleman et al., 2021). Here, too, two ensheathing and two ensheathing/wrapping glial cells are found in each hemineuromere (**Figure 2F,H**). Note, that dorsally to the neuropil, ensheathing glia send protrusions to the blood-brain barrier and thus adopt a more complex polarized morphology (**Figure 2H**). In consequence, dorsal neurons are encased by ensheathing glia processes and not by cortex glia (**Figure 1J**, **Figure 2H**, see below).

The light microscopic analysis suggests that ensheathing glial cells encase the neuropil in third instar larvae. This idea is supported by the ssTEM data sets. In L1 ventral nerve cord ensheathing, ensheathing/wrapping and astrocyte-like glial cells form an almost closed structure around the neuropil (**Figure 2C**). In contrast, in the L3 larval ventral nerve cord the neuropil encasement is complete (**Figure 2D**). In addition, multiple layers of glial processes are found around the neuropil (**Figure 2I,J**). At the ventral face of the neuropil, the glial sheath is often made of cell processes of ensheathing glial and cortex glial cells (**Figure 2I**).

### The complexity of ensheathing glia increases during development

About 112 ensheathing glial cells expressing *83E12-Gal4* can be detected in the third instar larva with only few ensheathing glial cells in the brain lobes (n=5, with 110, 112, 112, 114, and 116 *83E12-Gal4* positive cells, median = 112) **(Figure 3A, M)**, which corresponds to previously determined numbers (Beckervordersandforth et al., 2008; Kato et al., 2020; Omoto et al., 2015; Peco et al., 2016; Pereanu et al., 2005) and matches the annotations made in the TEM volume mentioned above. During the subsequent pupal development, the number of the *83E12-Gal4* positive ensheathing glial cell population increases more than 1,100 ensheathing glial cells in the adult (ventral nerve cord and brain without optic lobes, number determined using Imaris) **(Figure 3C,N)**.

**Figure 3.**
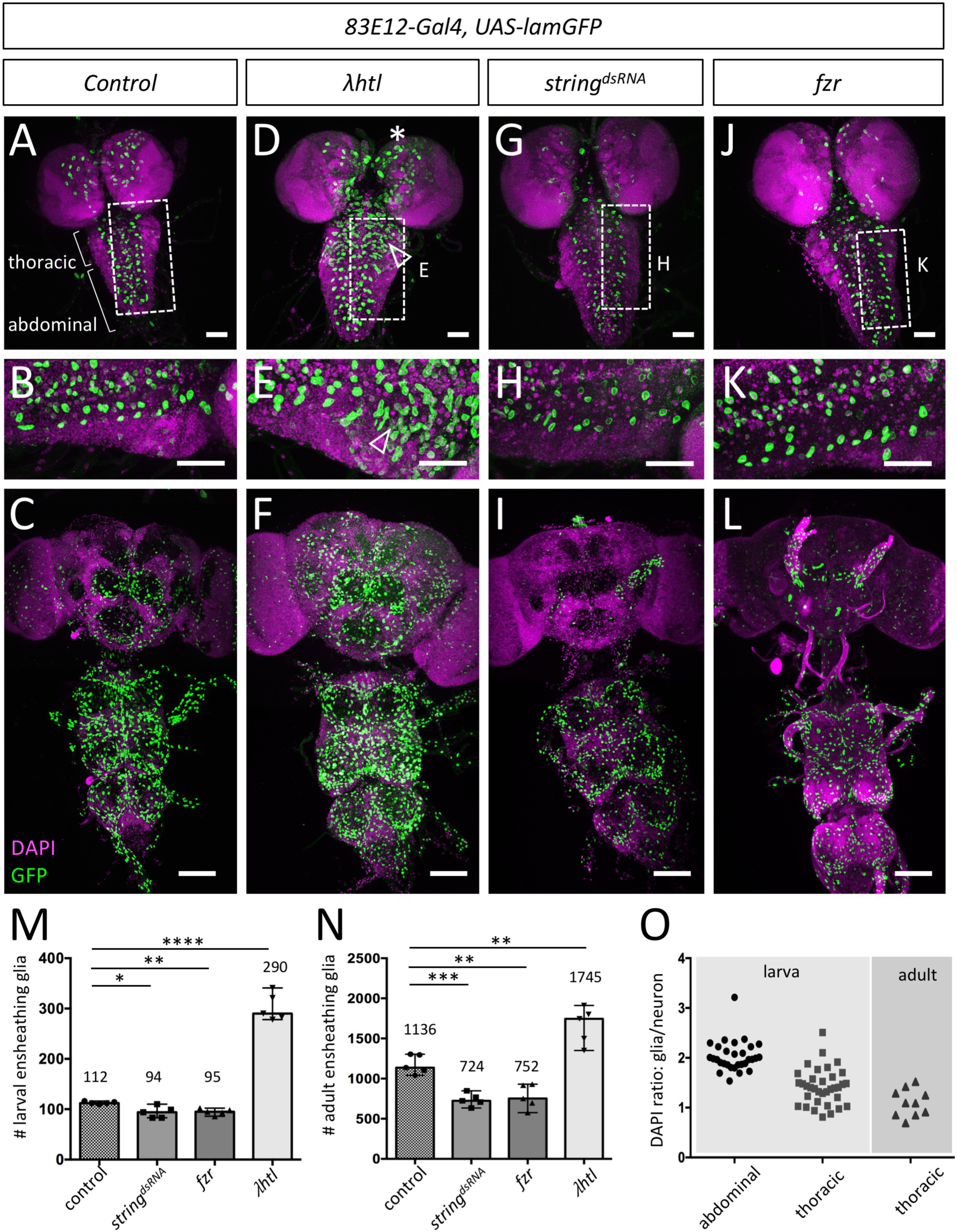
Larval thoracic ensheathing glial cells can divide to generate adult brain ensheathing glia (see also supplementary Figures 2-4) (A) Control third instar larvae with nuclei of ensheathing glia labelled [*83E12-Gal4 UAS-Lam::GFP*]. Nuclei are stained using DAPI. The boxed area is shown in high magnification in (B). (C) Adult control brain. The number of ensheathing glial nuclei is quantified in (M,N, n=5 brains for all genotypes). (D-F) Expression of an activated FGF-receptor leads to an increase in the number of larval ensheathing glia in larval thoracic neuromeres (arrowheads). The white boxed area is shown in higher magnification (E). (G-I) Upon expression of *string* dsRNA in all ensheathing glia the number of larval ensheathing glial cells is slightly reduced. For quantification see (M). The boxed area in the larval CNS (G) is shown in high magnification in (H). (I) Expression of *string* dsRNA in all ensheathing glia reduces the number of ensheathing glial cells in the adult CNS. (J-L) A similar reduction in the number of ensheathing glial cells is observed following expression of *fzr*. The boxed area in (J) is shown in high magnification in (K). Scale bars larval CNS (A,B,D,E,G,H,J,K) are 50 µm, scale bars for adult CNS (C,F,I,L) are 100 µm. Quantification of the ensheathing glial cell number in five larval brains (M) and in five adult brains (N) using Imaris (unpaired t-test). The optic lobes and the tract ensheathing glial cells were excluded in the quantification. For larva:; control – *string^dsRNA^*: * p = 0.0192; control – *fzr*: ** p = 0.0013; control - !htl: **** p =< 0.0001 for adult: *control* – *string^dsRNA^*: ***p = 0.0002; control – *fzr*: **p = 0.0018; control - *!htl*: ** p = 0.0057. (O) Quantification of DAPI intensity in 30 larval abdominal, 30 larval thoracic, and 10 adult thoracic ensheathing glial and neuronal nuclei.

It is unclear whether all larval ensheathing glia degenerate or whether some survive to participate in the formation of adult ensheathing glia (Kato et al., 2020; Omoto et al., 2015). To test whether ensheathing glial cells are competent to divide, we expressed activated FGF-receptor which is known to trigger perineurial glial proliferation (Avet-Rochex et al., 2012; Franzdóttir et al., 2009). Interestingly, a proliferative response upon activated FGF-receptor expression is most prominently observed in thoracic neuromeres (**Figure 3A-F,M,N, Figure S2**). We then blocked cell division in the ensheathing glia by RNAi-based silencing of *string*, which encodes the Drosophila Cdc25 phosphatase homolog required for progression from G2 to M phase (Edgar and O’Farrell, 1989). This expression regime did not severely affect ensheathing glia cell number in the larval CNS (**Figure 3G,H,M**, 94 ensheathing glia in *string* knockdown larvae vs. 112 ensheathing glial cells in control animals, n = 5 with 83, 83, 110, 99, and 94 *83E12-Gal4* positive cells, median = 94). In contrast, suppression of *string* function resulted in 38 % less ensheathing glial cells in the adult nervous system (724 [n=5, 724, 707, 769, 633, and 848 *83E12-Gal4* positive cells, median = 724] instead of the expected 1136 ensheathing glial cells **(Figure 3I,N)**. Similarly, when we expressed the cell cycle regulator Fizzy related (Fzr) which blocks cell division upon overexpression (Grosskortenhaus and Sprenger, 2002; Sigrist and Lehner, 1997; Silies and Klämbt, 2010) we noted a similar reduction in the number of adult ensheathing glia **(Figure 3J-N)**. Interestingly, the block of cell proliferation in ensheathing glia mostly affects the brain (**Figure 3I,J**). To further test whether larval ensheathing glia could divide, we analyzed the ploidy of the ensheathing glia using DAPI staining (Unhavaithaya and Orr-Weaver, 2012). In the larval ventral nerve cord, 30 of 30 tested abdominal glial nuclei are polyploid and thus are likely not to divide **(Figure 3O, Figure S3A-D)**. In contrast, 10 out of 25 ensheathing glial cells in thoracic neuromeres are diploid and possibly could divide **(Figure 3O, Figure S3A-D)**. Phospho-histone H3 serves as a specific marker for mitosis. In the larval CNS, we only rarely detected *83E12-Gal4* and phospho-histone H3 positive cells **(Figure S3E)**. Dividing ensheathing glial cells were easily detected in early pupae but not in 42 APF old pupae **(Figure S3F-H)**. In addition, we fed the thymidine analogue 5-ethynyl-2ʹ-deoxyuridine (EdU) which allows to label DNA synthesis during larval development and frequently detected EdU staining in *83E12-Gal4* positive ensheathing glia of larval and young pupal brains **(Figure S3I,J)**. Moreover, when we labelled larval ensheathing glia using a MCFO2 strategy (Nern et al., 2015) in third instar larvae or early pupae, we found labelled ensheathing glia in the adult (**Figure S4A-C**). In conclusion, at least some larval *83E12-Gal4* expressing ensheathing glial cells are diploid and can proliferate.

### Ensheathing glial cells are not required for viability

To test the functional relevance of the ensheathing glia, we performed ablation experiments. Expression of *hid* and *reaper* (*rpr*) using *repo-Gal4* or *83E12-Gal4* resulted in early larval lethality. Since *83E12-Gal4* is also expressed in the midgut, we used a split-Gal4 approach to restrict Gal4 expression to only the ensheathing glial cells of the CNS (Luan et al., 2006; Pfeiffer et al., 2010). We generated a construct that allowed the expression of a Gal4 activation domain under the control of the *83E12* enhancer [*83E12-Gal4^AD^*]. When crossed to flies that carry an element that directs expression of the Gal4 DNA binding domain in all glial cells [*repo-Gal4^DBD^*], Gal4 activity will be reconstituted only in CNS ensheathing glial cells. Indeed, animals carrying both transgenes [*83E12-Gal4^AD^, repo-Gal4^DBD^*] show the same expression domain as the original *83E12-Gal4* driver in the adult as well as in the larval nervous system (**Figure S4D-G)**.

Flies expressing *hid* and *rpr* only in the ensheathing glia [*83E12-Gal4^AD^*, *repo-Gal4^DBD^, UAS-hid UAS-rpr*] survive to adulthood. To validate the specificity of ensheathing glia ablation, we stained the specimens for Rumpel and Nazgul. Rumpel is a SLC5A transporter strongly expressed by the ensheathing glia and weakly by astrocyte-like glia **(Figure 4**, K. Yildirim personal communication). Nazgul is a NADP-retinol dehydrogenase that is specifically expressed by astrocyte-like cells (Ryglewski et al., 2017). In control, third instar larval brains Rumpel is found along the entire neuropil with Nazgul expressing astrocytes positioned in a characteristic pattern on the dorsal surface of the neuropil **(Figure 4A-C)**. Likewise, when we ablated the ensheathing glia in a background of a UAS-CD8::GFP transgene [*83E12-Gal4^AD^*, *repo-Gal4^DBD^, UAS-hid UAS-CD8::GFP*] no ensheathing glia cells can be detected in the CNS. Only few wrapping glial cells in the peripheral nervous system can still be detected **(Figure S4H-J)**.

**Figure 4.**
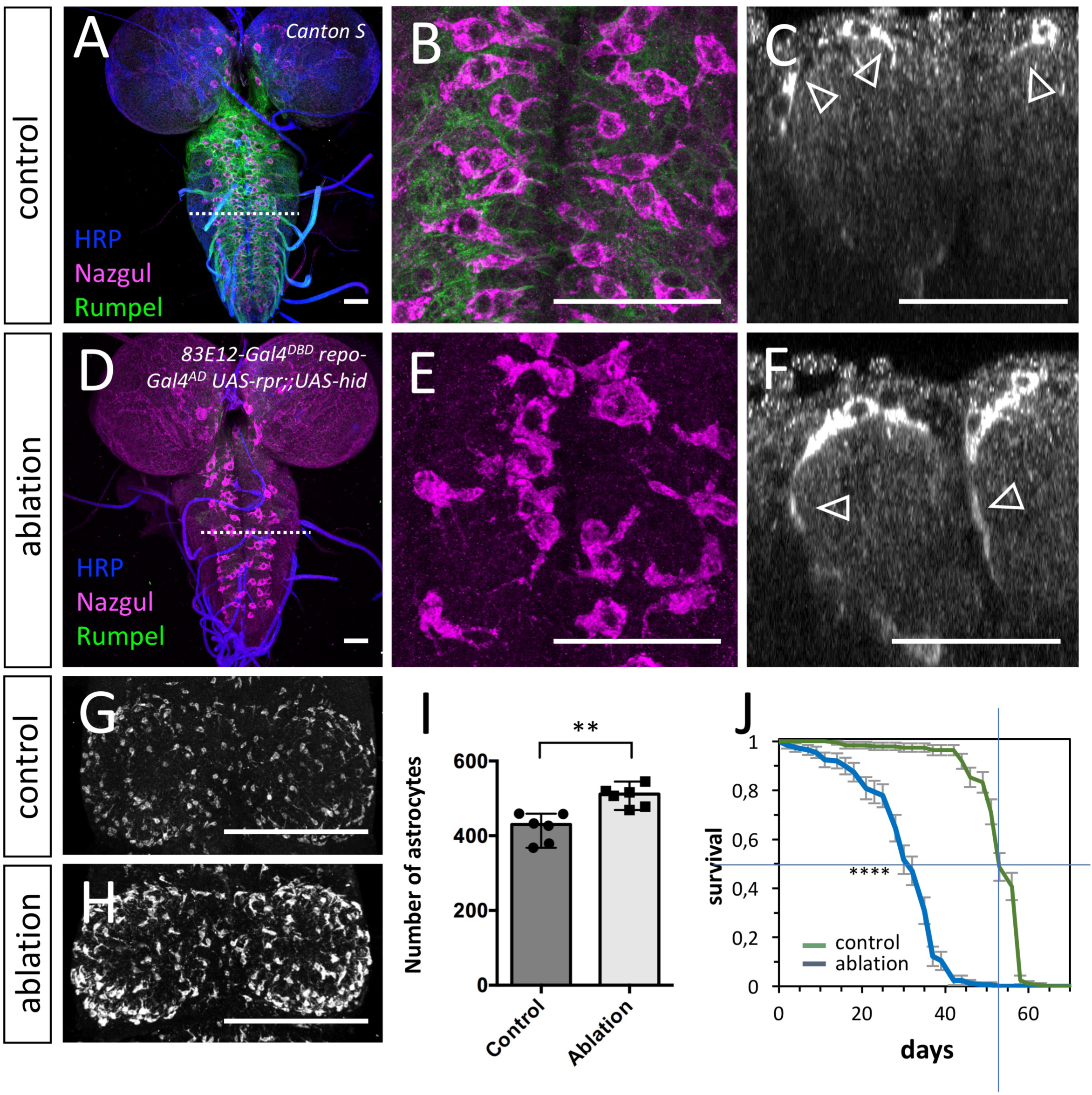
Ablation of ensheathing glia causes compensatory growth and increased proliferation of astrocyte-like cells (see also supplementary Figure 4) (A-F) Third instar larval brains stained for Rumpel (green, expressed by ensheathing glia), Nazgul (magenta, expressed by astrocyte-like cells) and HRP (blue, neuronal membrane marker). (A) Maximum projection of a confocal stack of a control larval CNS. The dashed line indicates the position of the orthogonal view shown in (C). (B) Dorsal view of a control ventral nerve cord. Note the regular positioning of the astrocyte-like cells (magenta). (C) Orthogonal section of astrocyte-like cells which are labelled using anti-Nazgul staining (white, arrowheads point to short protrusions). (D,E) Upon ablation of the ensheathing glia [*83E12-Gal4^DBD^ repo-Gal4^AD^ UAS-rpr;;UAS-hid*], Rumpel expression is greatly reduced. The dashed line indicates the position of the orthogonal view shown in (F). (E) Dorsal view of a larval ventral nerve cord following ensheathing glia ablation. The position and the morphology of the astrocyte-like cells appears disorganized. (F) Orthogonal section. Note, that astrocyte-like glial cells develop long cell processes that appear to encase the entire neuropil in the absence of ensheathing glia (arrowheads). (G) Dorsal view on a thoracic neuromere of an adult control fly. (H) Thoracic neuromere of an adult fly lacking ensheathing glial cells. The number of astrocyte-like cells increases. (I) Quantification of the number of astrocyte-like cells in the ventral nerve cord of adult flies (n = 6 brains, ** p = 0.0018, unpaired t-test). (J) Upon ablation of the ensheathing glia, longevity is reduced (n = 200 mated females, **** p = 4.67365E-83, Log-rank test). Scale bars are: (A-F): 50 µm, (G-H): 100 µm.

Astrocytes infiltrate the entire neuropil and form only few processes on the outer surface of the neuropil **(Figure 4C)**. Upon ablation of the ensheathing glia, only a very faint astrocytic Rumpel signal is detected **(Figure 4D,E)**. Astrocyte-like cells form additional large cell processes around the neuropil **(Figure 4F)**. In addition, in adult flies lacking ensheathing glia we noted a 20 % increase in the number of Nazgul positive astrocyte-like cells (**Figure 4G-I**, n=6 for each genotype, [control: 427, 459, 379, 458, 368 and 433, median = 430; ablation: 507, 520, 469, 478, 516, and 545, median = 512; ** p = 0.0022). Adult flies are fertile but lifespan is reduced by about 40 % compared to control animals (**Figure 4J**, n = 200 mated females, p = 4,67365E-83). In conclusion, these data show that ensheathing glial cells are not essential for viability. Upon ablation of ensheathing glia astrocyte-like cells form additional processes encasing the neuropil. This compensatory growth of astrocyte-like cells suggests that barrier establishment is a key function of the ensheathing glia.

### Ensheathing glial cells form a barrier around the neuropil

The third instar larval CNS is covered by subperineurial glial cells that do not allow penetration of labelled 10 kDa dextran. In wild type, subperineurial glial cells block paracellular diffusion across the blood-brain barrier by forming septate junctions (Stork et al., 2008). However, in larvae lacking septate junctions, extensive interdigitations between neighboring subperineurial cells can also provide a barrier function (Babatz et al., 2018).

Extensive electron microscopic analyses failed to demonstrate septate junction like structures between different ensheathing glial cells. However, glial processes overlap extensively at the neuropil cortex interface (**Figure 2I,J**), which might increase the length of the diffusion path at the boundary between CNS cortex and neuropil. To directly test whether ensheathing glial cells indeed provide a barrier function, we performed dextran diffusion assays in control brains and those lacking ensheathing glia. For this, we carefully dissected third instar larval brains including anterior cuticular structures and placed them on a coverslip. Upon injection of fluorescently labelled 10 kDa dextran directly into the neuropil of one brain lobe employing capillary normally used for DNA injections, we determined diffusion of the labelled dextran within the neuropil through the large brain commissure into the contralateral hemisphere **(Figure 5A)**. Diffusion of dextran was monitored under the confocal microscope for 10 minutes (**Figure 5B,D, movies S1,2**). The ratio of the fluorescence measured at two identical sized ROIs at the neuropil and the cortex contralateral to the injection site was plotted against time **(Figure 5C)**. In control larvae, a fluorescence is mostly confined to the neuropil area and only little dye reaches the cortex area. When we performed the injection experiments in larvae lacking ensheathing glia, we noted a significantly faster increase in fluorescence signal in the cortex area relative to the neuropil **(Figure 5C)**. Thus, we conclude that the ensheathing glia, although they lack specialized occluding junctions, indeed provide a barrier function possibly due to the extensive overlapping cell processes.

**Figure 5.**
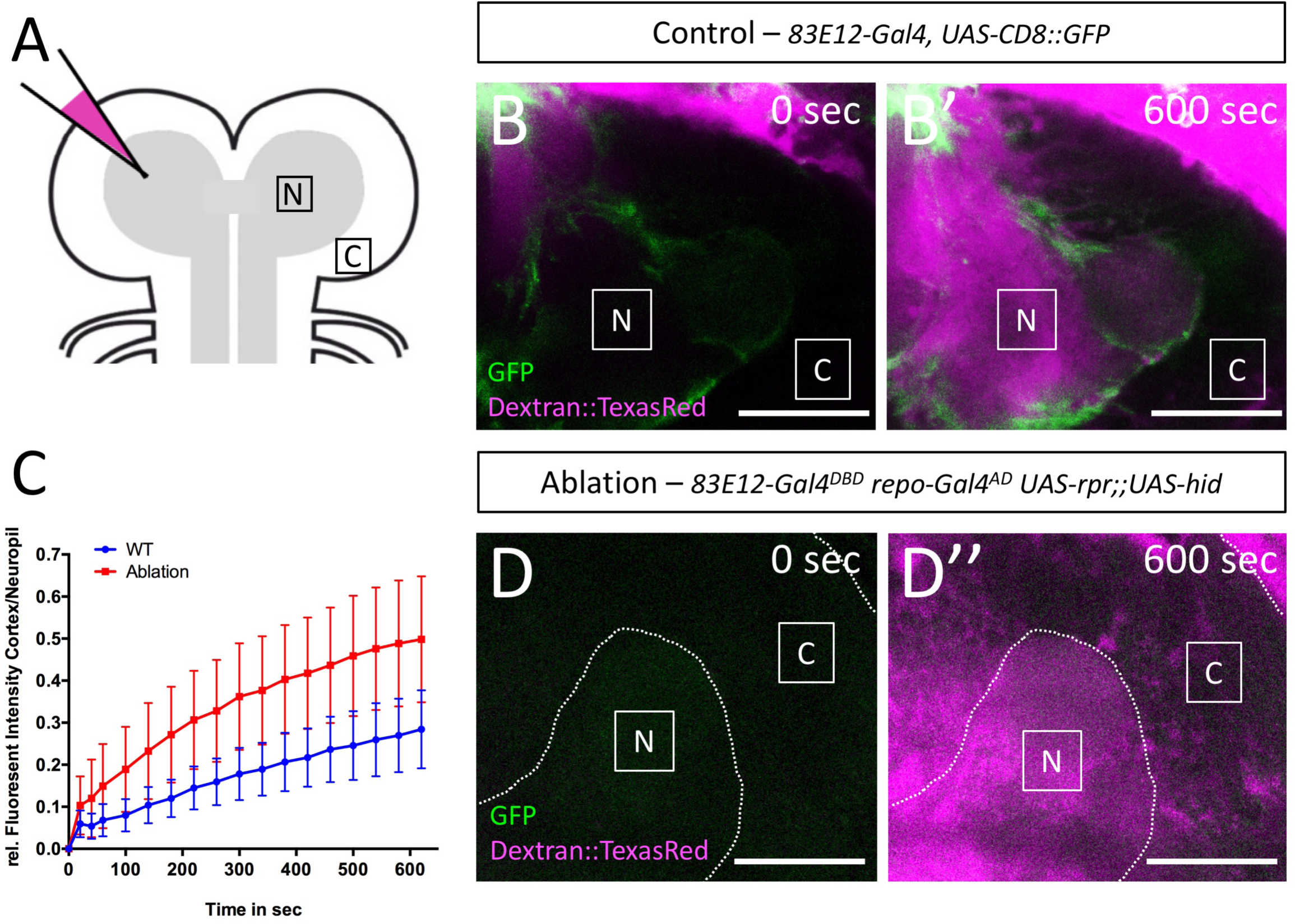
Ensheathing glial cells form an internal barrier around the neuropil (see also supplementary Movies 1,2) (A) Schematic depiction of the dye injection experiments. Fluorescently labelled 10 kDa dextran was injected into the left hemisphere of a wandering third instar larva. Diffusion of the dye was monitored at two positions. In the neuropil (N) and in the cortex (C) harboring all neuronal cell bodies. (B-B’) Stills of a movie of a larval brain lobe with the genotype [*83E12 UAS-CD8::GFP*] following dye injection, time is indicated. (C) Quantification of dye diffusion rate in larvae of the genotype [*83E12 UAS-CD8::GFP*] (blue line) and [*83E12 UAS-CD8::GFP, UAS-rpr;;UAS-hid*] (red line) (n=5, standard deviation is shown, *p = 0.0181; unpaired t-test). (D-D’) Stills of a movie of a larval brain lobe with the genotype [*83E12 UAS-CD8::GFP, UAS-rpr;;UAS-hid*] following dye injection. Scale bars are 50 µm.

### Ensheathing glial cells are polarized

Tissue barriers are generally formed by polarized cells which are characterized by a differential distribution of plasma membrane lipids. The apical membrane is generally rich in phosphatidylinositol 4,5-bisphosphate (PIP_2_), whereas the basolateral cell membrane contains more phosphatidylinositol 3,4,5-triphosphate (PIP_3_) (Krahn, 2020; Shewan et al., 2011). To detect a possible differential localization of these phospholipids, we employed different lipid sensors based on distinct PH-domains. PH-PLCδ-mCherry is targeted by PIP_2_ (Khuong et al., 2010) whereas PH-AKT-GFP preferentially binds PIP_3_ (Ivetac et al., 2009). We focused our analysis on the ensheathing glial cells covering the dorsal aspect of the neuropil.

We co-expressed both sensors in the ensheathing glia using *83E12-Gal4* and quantified the distribution of both sensors by determining the number of green and red fluorescent pixels in *83E12-Gal4* positive cells in orthogonal sections comparing the cell domain close to the neuropil with the one close to the blood-brain barrier (**Figure 6A-G**). Here, we noted an increased PH-PLCδ-mCherry localization at the direct interface of the neuropil and the ensheathing glia (**Figure 6B-E**). For quantification, we measured the mean fluorescence intensity of PH-PLCδ-mCherry and PH-AKT-GFP (30 dorsal neuropil areas of 10 larvae). This demonstrated an almost twofold enrichment of PIP_2_ at the plasma membrane domain facing the neuropil. The PIP_3_ detecting PH-AKT-GFP shows a complimentary distribution with an enrichment in basolateral plasma membrane domains **(Figure 6F,G)**. A similar polarization of the ensheathing glial cells was detected in adult stages (**Figure 6F**; **Figure S5**, 30 neuropil areas in 10 adult brains).

**Figure 6.**
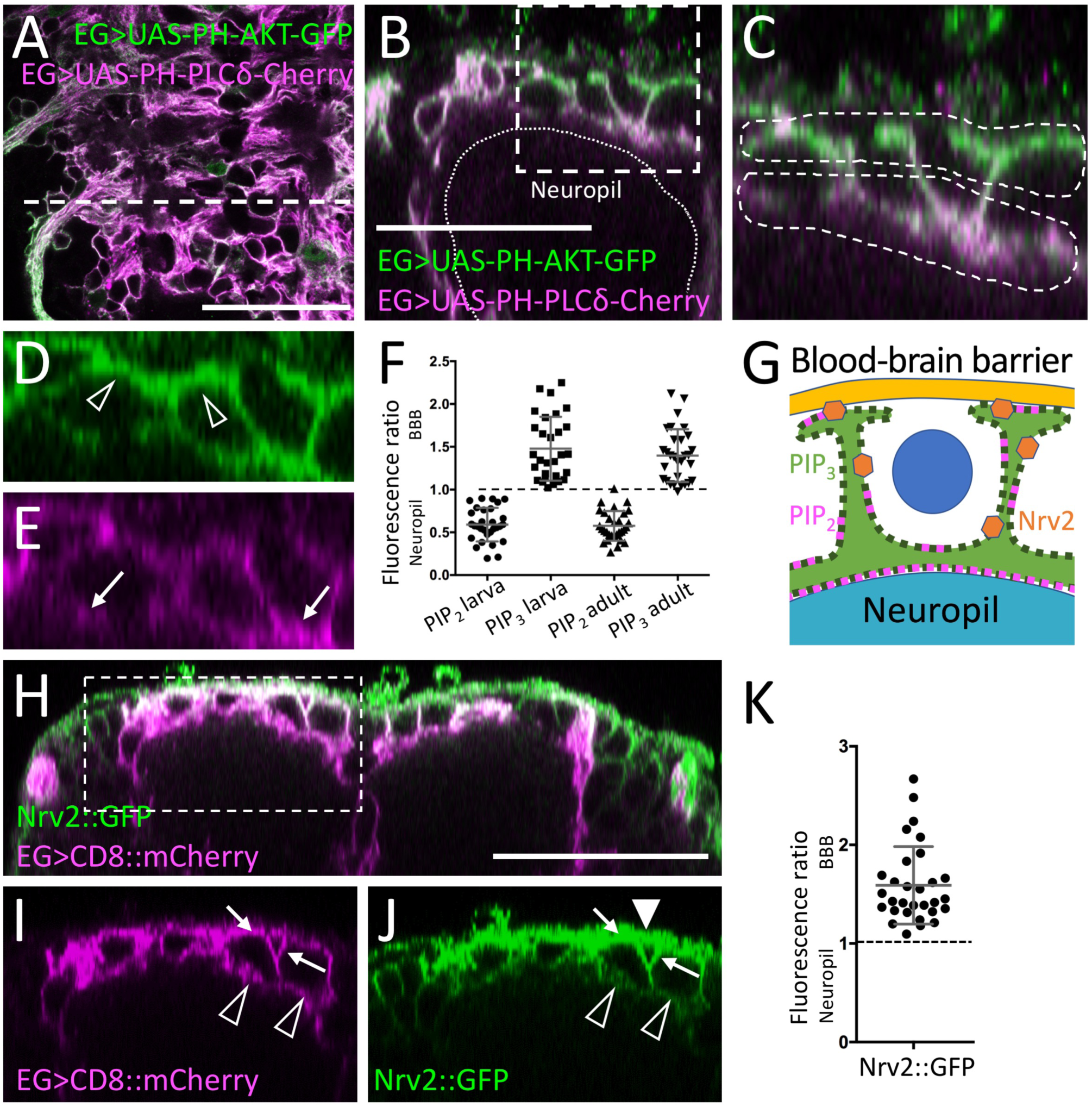
Ensheathing glial cells show polarized plasma membrane domains (see also supplementary Figure 5) (A) Projection of a confocal stack of a third instar larval ventral nerve cord coexpressing *PH-AKT-GFP* and *PH-PLCδ-mCherry* in ensheathing glial cells (EG) [*83E12-Gal4, UAS-PH-AKT-GFP, UAS*-*PH-PLCδ-mCherry*]. The dashed white line indicates the position of the orthogonal section shown in (B). (B) Orthogonal section showing the dorsal aspect of the neuropil. The boxed area is shown in higher magnification in (C-E). The dashed areas were subsequently used for quantification of GFP and mCherry localization. (F) Quantification of GFP/mCherry distribution in larval and adult ensheathing glia. The ratio of red (PLCδ-mCherry shown in magenta) and green (PH-AKT-GFP) fluorescent pixels of the areas indicated in (C) is plotted. (G) Schematic view of a dorsal ensheathing glial cell. The neuropil facing domain is characterized by a high PIP_2_ content, whereas PIP_3_ is concentrated on the baso-lateral domain, where Nrv2 is predominantly localized, too. The blue dot indicates the position of a neuron. (H) Coexpression of *Nrv2::GFP* and *83E12-Gal4, UAS-mCherry* in the ventral nerve cord of third instar larva. The boxed area is shown in higher magnification in (I,J). Note the preferential localization of Nrv2 at the basolateral cell domain of the ensheathing glia (arrows). Only little Nrv2 is found at the apical domain (open arrow head). Additional expression of Nrv2 is seen in the blood-brain barrier (filled arrowhead). (K) Quantification of polarized Nrv2::GFP localization. Scale bars are for larva 50 µm.

Polarized epithelial cells are characterized by an enrichment of the Na^+^ / K^+^-ATPase Nervana 2 (Nrv2) at the basolateral plasma membrane (Dubreuil et al., 2000). To study the localization of Nrv2, we employed a gene trap insertion (Morin et al., 2001). As found for PIP_3_, Nrv2^GFP^ localizes predominantly at the basal side of the ensheathing glia **(Figure 6H-K)**, which corresponds to the localization of Nrv2 in epithelial cells. Therefore, we conclude that ensheathing glial cells are polarized cells throughout development, facing their apical-like domain towards the neuropil (**Figure 6G**).

### Integrin α and extracellular matrix components are expressed by adult ensheathing glia

In epithelial cells, apical-basal polarity is also reflected by a basally located integrin receptor which bind the basally located extracellular matrix (ECM) proteins. *inflated* encodes an alpha-subunit of the integrin receptor and *inflated* mRNA is expressed by adult perineurial and ensheathing glial cells (Bökel and Brown, 2002; Davie et al., 2018) **(Figure S6)**. Endogenously YFP-tagged Integrin *α (if^CPTI-004152^*, (Lowe et al., 2014)) localizes at the blood-brain barrier and around the neuropil in larval brains **(Figure 7A,B,E)**. In the adult brain, Integrin^YFP^ *α* localization around the neuropil appears more pronounced compared to abdominal neuromeres **(Figure 7C,D)**. Integrin receptors bind ECM components, suggesting that these proteins are also expressed within the nervous system. We thus tested expression of all genes annotated to encode proteins involved in extracellular matrix formation (FlyBase) using published single cell sequencing data of adult brains (Davie et al., 2018) **(Figure S6)**. Eight of these genes appear expressed by the ensheathing glia and GFP-protein trap insertion lines were available for *trol* (Perlecan), *viking* (Collagen IV), and *dally* (Glypican).

**Figure 7.**
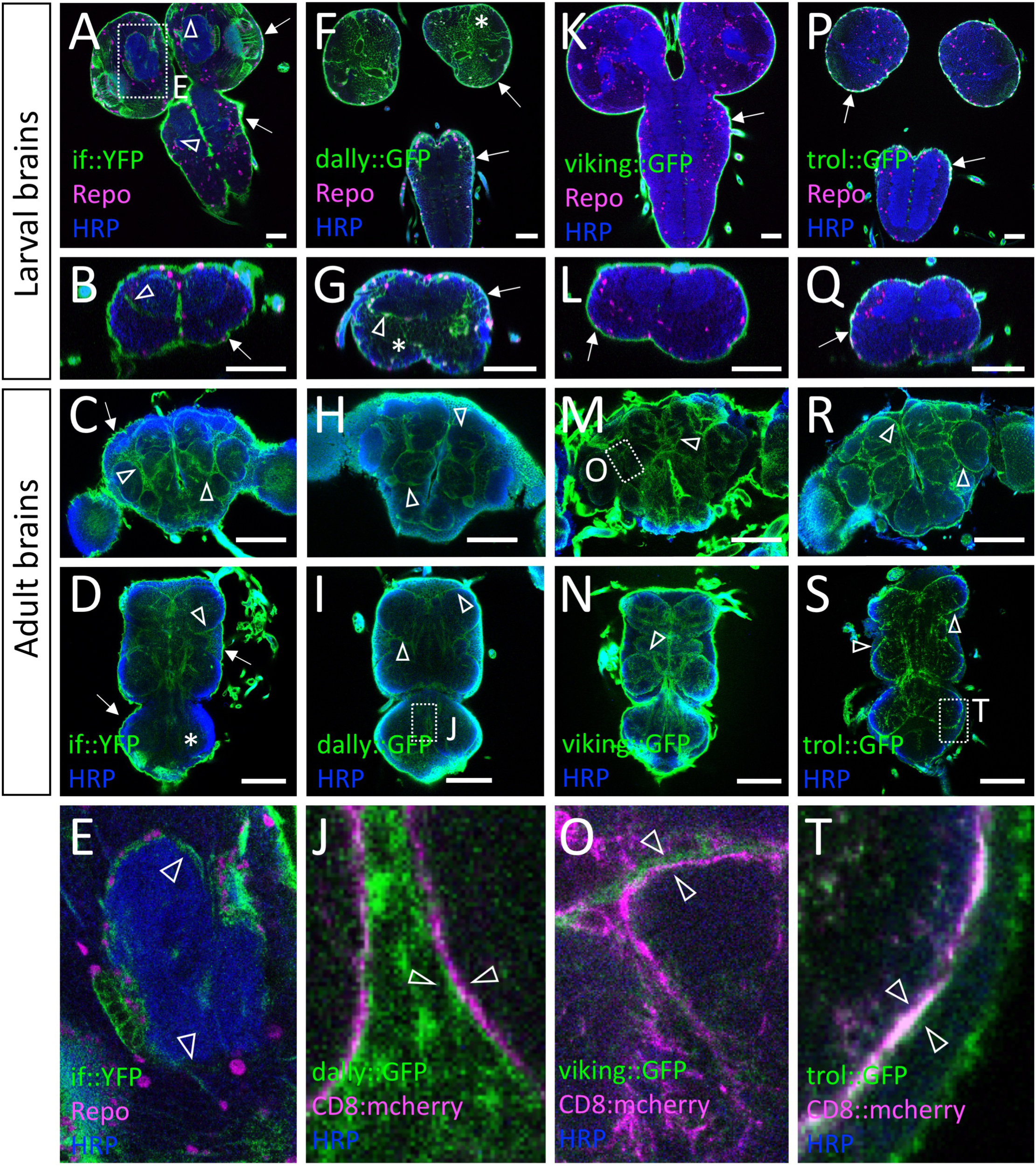
The ensheathing glial cells are flanked by extracellular matrix (see also supplementary Figure 6) Expression of Integrin α and different extracellular matrix components as detected by using gene trap insertion lines. (A-E) Endogenously YFP-tagged Integrin α encoded by the *inflated* gene. (F-J) Endogenously GFP-tagged heparan sulfate proteoglycan Dally, (K-O), the Collagen IV protein encoded by *viking*, and (P-T) the Perlecan protein encoded by *trol*. The different developmental stages are indicated. (A,I,M,S) The boxed areas are shown in the bottom row. Glial cells are in magenta (anti-Repo staining), neuronal membranes are in blue (anti-HRP staining) and the GFP-tagged proteins are in green (anti-GFP staining). (A,B,E) During late larval stages, prominent Inflated expression is seen at the blood-brain barrier (arrows) and weak expression is seen around the neuropil at the position of the ensheathing glia (arrowhead). (C,D) In adults, Integrin α is still found at the blood-brain barrier (arrows) and becomes more prominent around the neuropil (arrowheads). Note that strongest Inflated expression is detected in larval brains. No Inflated expression can be detected in the abdominal neuromeres (asterisk). (F,G) During the larval stage Dally is expressed at the blood-brain barrier (arrows) and in the CNS cortex (asterisks). A slightly stronger localization of Dally can be detected at the position of basal ensheathing glia processes (arrowhead). (H-J) In adults, Dally is enriched at the position of the ensheathing glia (arrowheads). (K,L) In the larval CNS Collagen IV is found at blood-brain barrier (arrows) but not within the nervous system. (M-O) Collagen IV is detected at the neuropil-cortex interface in adult stages (arrowheads in M,N, boxed area is shown in O). (P,Q) Trol is detected at the larval blood-brain barrier (arrows) but not around the neuropil. (R-T) In adults, prominent Trol localization is seen at the position of the ensheathing glia (arrowheads). The boxed area in (S) is shown at higher magnification in (T). Scale bars are: larval CNS 50 µm, adult CNS 100 µm.

The heparan sulfate proteoglycan Dally is most strongly detected close to the cells of the larval blood-brain barrier **(Figure 7F)**. Within the larval CNS, Dally is found in the cortex and is enriched around the larval neuropil **(Figure 7F,G)**. In adults, Dally localization at the blood-brain barrier ceases but is still found in the CNS cortex. A prominent enrichment of Dally is seen around the neuropil **(Figure 7H-J)**. In contrast to Dally, we could not detect Trol and Viking around the larval neuropil and only detected strong signals at the blood-brain barrier **(Figure 7K,L,P,Q)**. However, in the adult nervous system both Trol and Viking are detected basally of the ensheathing glia facing the cortex glia with an even distribution in the different parts of the CNS **(Figure 7M-O,R-T)**. In conclusion, the ensheathing glia express ECM proteins which is characteristic for polarized cell types.

### Polar distribution of ß_Heavy_-Spectrin in larval ensheathing glia

Polarization of cells is also evident in the cytoskeleton underlying the plasma membrane. Whereas ⍺-Spectrin is found below the entire plasma membrane, ß-Spectrin decorates the basolateral plasma membrane and ß_Heavy_-Spectrin (ß_H_-Spectrin) is located at the apical domain of the cell (Thomas and Kiehart, 1994). ß-Spectrin is found in all neural cells in the ventral nerve cord (Hülsmeier et al., 2007) and no specific localization can be resolved in the ensheathing glia due to its strong overall expression. ß_H_-Spectrin is encoded by the *karst* gene. One available MiMIC insertion allows to target two of the seven isoforms **(Figure 8A)**. In third instar larval brains, GFP-ß_H_-Spectrin localizes at the blood-brain barrier **(Figure 8C,D,F,G)**, and close to the neuropil **(Figure 8C,D,F)**, indicating expression by the ensheathing glia. Here, ß_H_-Spectrin is enriched close to the apical-like, PIP_2_ containing membrane of the ensheathing glia facing towards the neuropil **(Figure 8C,D,K)**. To further validate the expression of Karst in the ensheathing glia, we silenced *karst* expression by expressing double stranded RNA. When *karst* expression is silenced in all glial cells using *repo-Gal4* only expression in trachea as well as a weak background signal at the neural lamella around the CNS is detected (**Figure S7A,B**). Silencing of *karst* expression in the blood-brain barrier using *moody-Gal4* did not affect localization of GFP around the neuropil (**Figure S7C,D**). When we silenced *karst* specifically in the ensheathing glia using *83E12-Gal4* protein localization in the blood-brain barrier is unaffected but no ß_H_-Spectrin protein can be detected around the neuropil **(Figure 8G)**.

**Figure 8.**
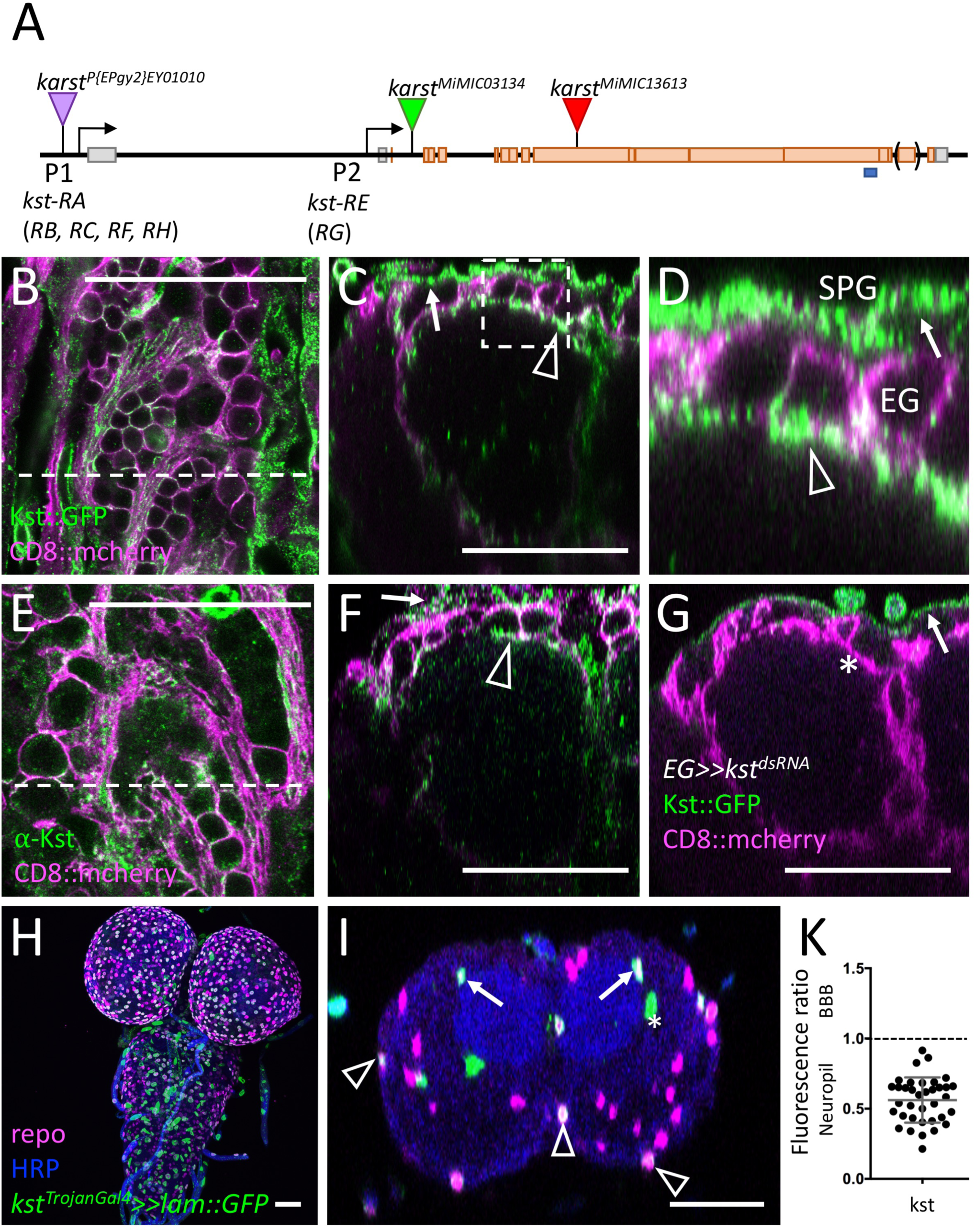
ß_H_-Spectrin shows a polar distribution in ensheathing glia (see also supplementary Figures 8,7) (A) Schematic view of the *karst* locus. Transcription is from left to right. Seven different karst mRNAs are generated from two promoters (P1, P2) as indicated (*kst-RA - RH*). The ß_H_-Spectrin proteins PE and PG differ in a C-terminal exon indicated by brackets. The position of two MiMIC insertions and the EP insertion used for gain of function experiments is indicated, the blues line denotes the position of the peptide used for immunization. (B) Confocal view of the surface of a third larval instar brain of a *kst^MiMIC03134::GFP^* animal that expresses mCherry in all ensheathing glia [*83E12-Gal4, UAS-CD8::mCherry*]. The dashed line indicates the position of the orthogonal view shown in (C,D). Expression at the blood-brain barrier forming subperineurial glia (arrows) and at the apical domain of the ensheathing glia (arrowheads) can be detected. For quantification see (K). (E) Similar view as shown in (B) of a control larva stained for Kst expression using a polyclonal antiserum. The dashed line indicates the position of the orthogonal view shown in (F). (F) Localization of Kst protein is weakly seen in the blood-brain barrier (arrow) and the ensheathing glia (arrowhead). (G) Orthogonal view of a *kst^MiMIC03134::GFP^* animal expressing *kst^dsRNA^* in the ensheathing glia (EG) [*83E12-Gal4, UAS-kst^dsRNA^*]. Expression of Karst in ensheathing glia cannot be detected anymore (asterisk). Expression in the blood-brain barrier is unaffected (arrow). (H,I) The Trojan Gal4 element in *kst^MiMINC03134^* directs Gal4 expression in the *kst P2* pattern. Lamin::GFP localizes to ensheathing glial nuclei (arrows), some cells of the blood-brain barrier (arrowheads) and some cells in the position of tracheal cells (asterisks). (K) Quantification of Karst localization in different membrane domains of ensheathing glial cells, quantification as described in Figure 6. Scale bar is 50µm.

A similar ß_H_-Spectrin localization was detected using newly generated antisera directed against a C-terminal peptide present in all predicted ß_H_-Spectrin isoforms **(Figure 8A,E,F**, for specificity of the generated antiserum see: **Figure S7E,F)**. Furthermore, we generated a Trojan-Gal4 insertion using the same MiMIC insertion that was used to generate the Karst^GFP^ protein trap **(Figure 8A)**. The *kst^Trojan-Gal4^* allele is expected to report only the expression pattern of the proximal promoter which activates expression of two out of the seven *karst* splice variants (FlyBase) **(Figure 8A)**. Indeed, when driving a nuclear GFP reporter (*UAS-Lam::GFP*) that allows labelling of nuclei, expression in the ensheathing and blood-brain barrier glial cells is apparent **(Figure 8H,I)**. In addition, cells in the position of tracheal cells express the *kst^Trojan-Gal4^* allele **(Figure 8I)**. During pupal stages, expression of Karst ceases between 72 and 96 hours after puparium formation (APF) (**Figure S8**). Adult ensheathing glial cells lack detectable expression of the *kst^MiMIC::GFP^* insertion and show no reactivity using the anti-Karst antisera.

### ß_H_-Spectrin is required for ensheathing glial polarity

We next tested whether ß_H_-Spectrin is required for ensheathing glial cell polarity and determined the overall morphology of the ensheathing glia in the *karst* loss of function *karst^MiMIC13613^* larvae using *83E12-Gal4* driving *CD8::GFP*. In *karst* mutants, larval ensheathing glial cells are still present but show an abnormal collapsed morphology. The extension of ensheathing glia cell protrusions around dorsally located neuronal cell bodies is less evident **(Figure 9A,B)**. When we silenced *karst* expression using RNAi, we noted a weaker phenotype and dorsal protrusions frequently formed **(Figure 9C)**. This allowed an analysis of the distribution of PIP_2_ and PIP_3_ **(Figure 9C-E)**. Whereas in ensheathing glial cells of control ventral nerve cords, PIP_2_ (as detected by PH-PLCδ-mCherry) is found predominantly on the domain facing the neuropil, an almost even distribution is noted upon *kst* knockdown in larvae (compare **Figure 6F**, **Figure 9I**). Interestingly, the polar distribution of PIP_3_ does not appear to be affected by *kst* knockdown **(Figure 9E,I)**. To further test whether ß_H_-Spectrin is needed for ensheathing glial polarization, we performed overexpression experiments. To direct expression of wild type Karst protein in ensheathing glia, we utilized the EP-element insertion *P{EPgy2}EY01010*. Overexpression of *karst* results in abnormally convoluted and partially collapsed ensheathing glial cell morphology characterized by an even distribution of both PIP_2_ and PIP_3_ (**Figure 9F-H,J**).

**Figure 9.**
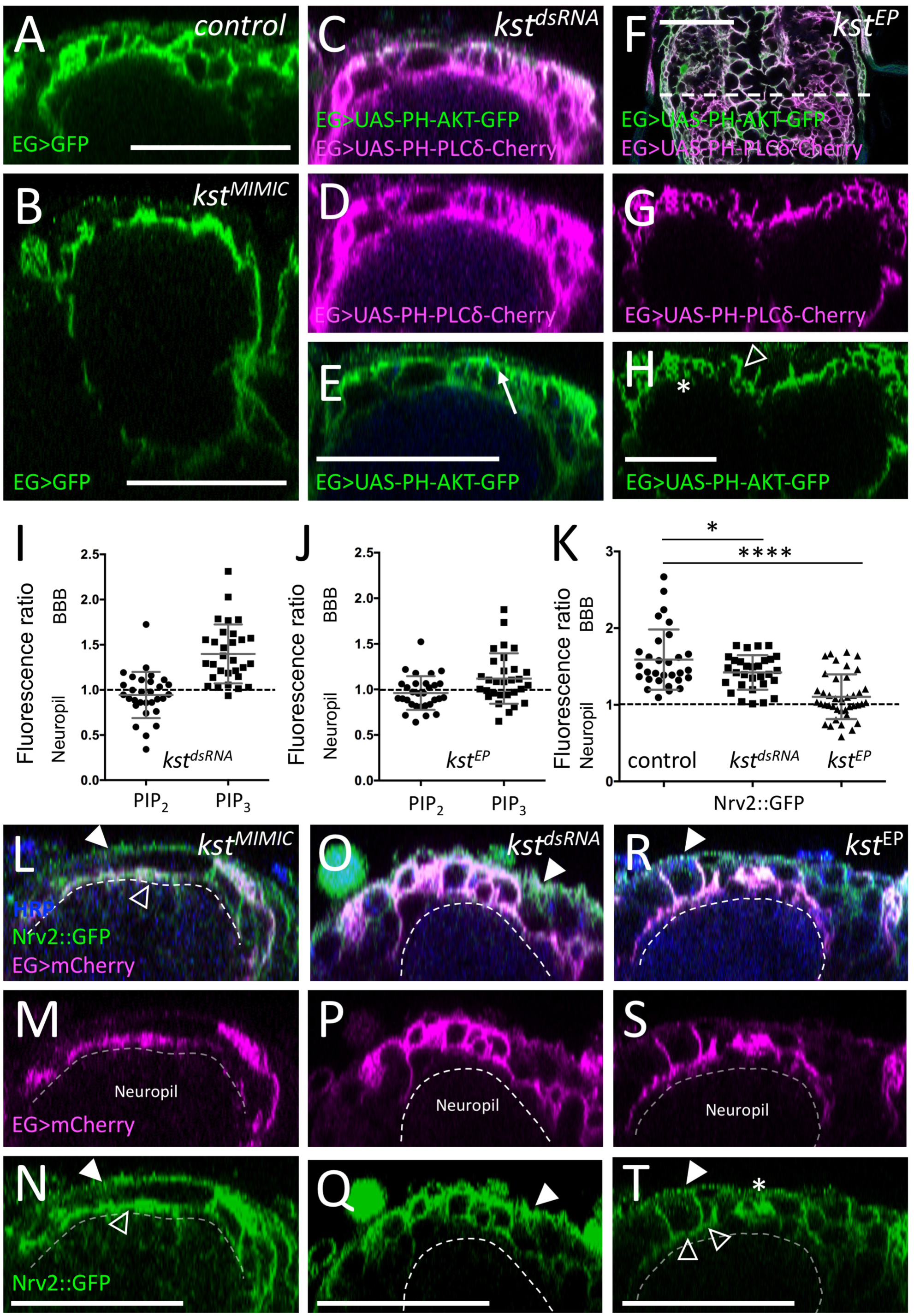
The ß_H_-Spectrin cytoskeleton is required for glial polarity. (A) Orthogonal view of the dorsal aspect of a third instar control ventral nerve cord stained for ensheathing glial (EG) morphology [*83E12-Gal4 UAS-CD8::GFP*]. (B) Third instar larval brain of a zygotic *karst* null mutant larvae with the genotype [*kst^MIC13613^ / kst^MIC13613^, 83E12-Gal4 UAS-CD8::GFP*]. The absence of all ß_H_-Spectrin protein affects the ensheathing/wrapping glial cell morphology. (C-E) Third instar larval ventral nerve cord with reduced *kst* expression in ensheathing glia [*83E12-Gal4 UAS-kst^dsRNA^, UAS-PH-AKT-GFP, UAS-PH-PLCδ-Cherry*]. (D) PH-PLCδ-Cherry binds to PIP_2_ and shows an even distribution between apical and basolateral plasma membrane domains. For quantification see (I). Quantification was performed as described in Figure 6. (E) PH-AKT-GFP binds to PIP_3_ and is distributed in a polarized manner. For quantification see (I). (F-H) Third instar larval ventral nerve cord with increased *kst* expression in ensheathing glia [*83E12-Gal4, kst^P{EPgy2}EY01010^, UAS-PH-AKT-GFP, UAS-PH-PLCδ-Cherry*]. Note, the variable phenotype noted upon *karst* overexpression. Ensheathing glia with almost normal morphology (open arrowhead) is next to hyperconvoluted ensheathing glia (asterisk). Quantification of PIP_2_ and PIP_3_ localization is shown in (J). (I,J) Quantification was performed as described in Figure 6. (L-N) In *karst* mutant larvae, Nrv2 is found in ensheathing glia cell processes (open arrowhead) that face the neuropil. The dashed line indicates the neuropil ensheathing glia boundary. (O-Q) Upon ensheathing glial specific *kst* knockdown, Nrv2 localizes to the basolateral cell domains, for quantification see (K). Filled arrowhead indicates the blood-brain barrier. (R-T) Upon overexpression of Karst, polarized localization of Nrv2 is less apparent in ensheathing glial cell processes (normal morphology: open arrowheads, hyperconvoluted morphology: asterisk), for quantification see (K). Scale bars are 50µm.

We next assayed Nrv2 localization in *karst* null mutants as well as upon *83E12-Gal4* directed *kst* knockdown. In larvae lacking *karst* expression ensheathing glial cells show a collapsed morphology and polarized localization of Nrv2 cannot be resolved **(Figure 9K-M)**. Upon *kst* knockdown Nrv2 correctly localizes to basolateral protrusions **(Figure 9K, O-Q)**. In *kst* gain of function larvae, the polarized Nrv2 distribution is lost **(Figure 9K,R-T)**, supporting the notion that the spectrin cytoskeleton might affect proper positioning of Nrv2 (Dubreuil, 2006; Dubreuil et al., 2000; Knust, 2000; Liem, 2016).

### Polarized ensheathing glia is required for locomotor behavior

The above data show that larval ensheathing glia are polarized, ECM abutting cells that separate the neuropil from the CNS cortex and are required for longevity of the adult fly. To test whether the ensheathing glial cells are required for normal locomotor control during larval stages, we compared locomotion of control animals with those lacking ensheathing glia or with those with reduced ß_H_-Spectrin or Nrv2 expression using FIM imaging (Risse et al., 2017; 2013).

Control animals move on long paths interrupted by short reorientation phases that are characterized by increased body bending (**Figure 10A**). *kst* knockdown specifically in ensheathing glial cells [*83E12-Gal4^AD^, repo-Gal4^DBD^, UAS-kst^dsRNA^*] causes a reduction in the peristalsis efficiency during go-phases **(Figure 10A,B,F,G)**. Likewise, crawling velocity is reduced significantly **(Figure 10G)**. This suggests that the specific lack of ß_H_-Spectrin in ensheathing glia causes a strong locomotor phenotype. We next analyzed mutant larvae to further validate the RNAi-induced phenotype. The Trojan-Gal4 insertion in the *karst^MiMIC03134^* insertion is expected to affect only isoforms Karst-PE and Karst-PG **(Figure 8)**. As control, we used an insertion of the Gal4 element in the opposite, unproductive orientation. Similar as detected for the *kst* knockdown, we noted a decreased peristalsis efficiency and a reduced crawling velocity **(Figure 10 C,I,J)**. Larvae completely lacking zygotic *karst* expression [*karst^MiMIC13613^* / *Df(3L)ED2083*] show a comparable larval locomotion phenotype (**Figure 10D,I,J**). This larval locomotion phenotype is similar to the one observed following ablation of the ensheathing glial cells using the genotype [*83E12-Gal4^AD^, repo-Gal4^DBD^, UAS-hid, UAS-rpr*] (**Figure 10E**). Thus, we conclude that polarized ensheathing glia that connect the blood-brain barrier with the dorsal neuropil are required for normal locomotor behavior. In order to perform vectorial transport, the Na^+^/K^+^ ATPase must act in a polarized fashion. We thus also compared larval locomotion of animals with reduced *nrv2* expression to control animals. Interestingly, larvae with ensheathing glial cells lacking *nrv2* expression behave opposite to larvae that lack *ß_H_-spectrin* showing an increased peristalsis efficiency as well as an increased crawling velocity **(Figure S9)**. The analysis of the role of polarized Nrv2 distribution for ensheathing glia physiology will thus be an interesting topic for future research.

**Figure 10.**
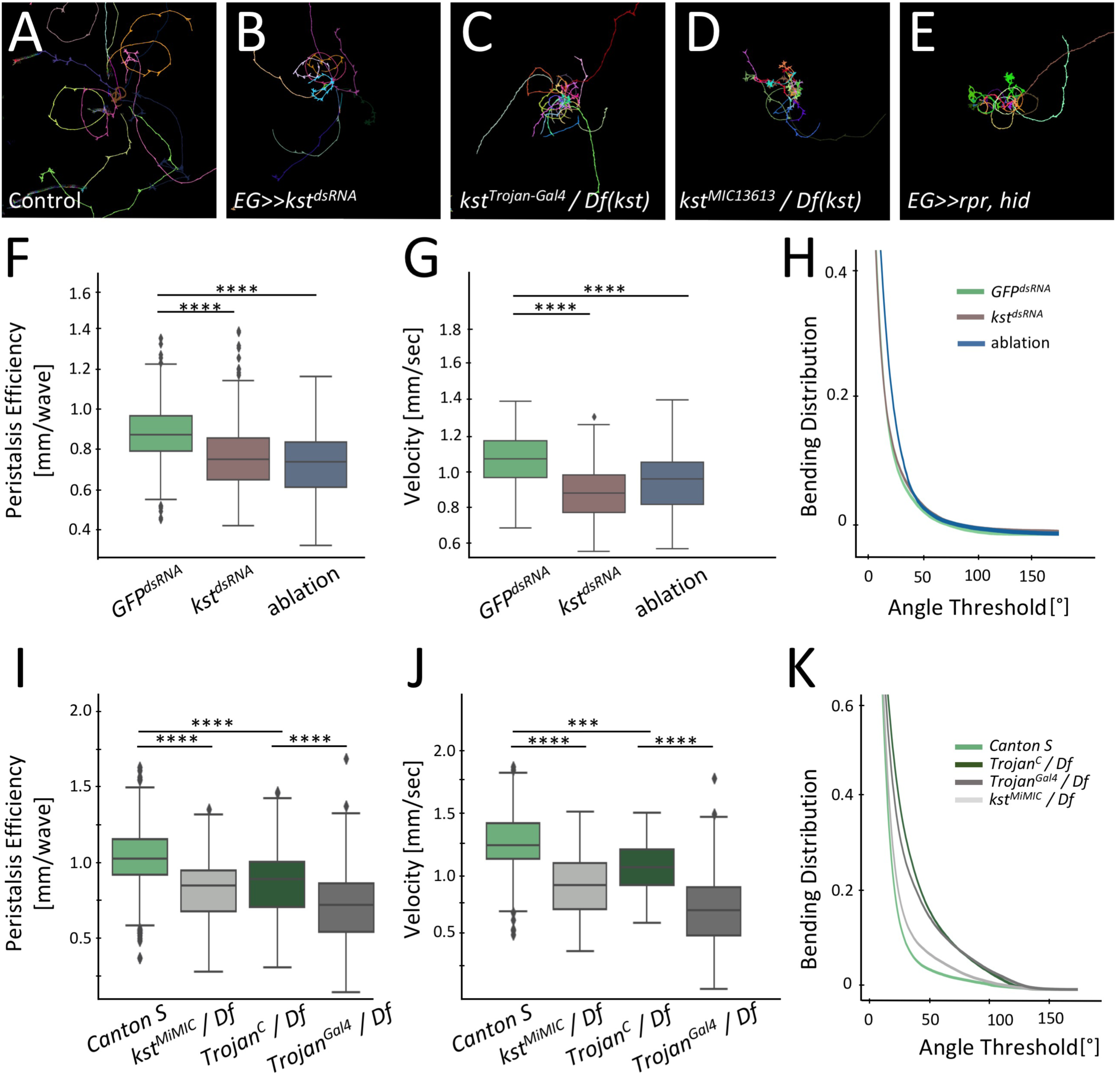
Karst is required in ensheathing glia for normal locomotor behavior (see also supplementary Figure 9) (A) Exemplary locomotion tracks of control wandering third instar larvae with the genotype [*83E12-Gal4^AD^, repo-Gal4^DBD^, UAS-GFP^dsRNA^*]. Larvae move in long run phases. (B) Upon *kst* knockdown in ensheathing glia (EG) only [*83E12-Gal4^AD^, repo-Gal4^DBD^, UAS-kst^dsRNA^*], larvae do not explore the tracking arena. (C) The insertion of a Trojan Gal4 element into the first coding exon of *kst-RE(RG)* generates an PE and PG isoform specific mutant (see Figure 8). Larvae with the genotype [*kst^TrojanGal4^ / Df(3L)ED208*] show a comparable locomotion phenotype as RNAi knockdown larvae. (D) Zygotic *karst* null mutant larvae with the genotype [*kst^MIC13613^ / Df(3L)ED208*] show a locomotion phenotype that is comparable to what we observe following specific ablation of the ensheathing glia ([*UAS-rpr, 83E12-Gal4^AD^, repo-Gal4^DBD^, UAS-hid*], E). (F-H) Quantification of peristalsis efficiency, velocity and bending distribution between control larvae, *kst* knockdown and ensheathing glia ablated animals. Box plots show median (line), boxes represent the first and third percentiles, whiskers show standard deviation, diamonds indicate outliers. (I-K) Quantification of the same parameters comparing wild type Canton S with *kst* null larvae and a *kst^Trojan^* control strain that carries the Trojan insertion in a non-productive orientation [*kst^MiMC03134-Trojan-C^* in trans to the *Df(3L) ED208*] with the isoform specific *kst^PE, PG^* mutant [*kst^MiMINC03134-TrojanGal4^ / Df(3L)* ED208. Quantification Wilcoxon rank-sum test, n=50. Peristalsis Efficiency [mm/wave]: *GFP^dsRNA^* – kst^dsRNA^: ****p = 2.65E-32 *GFP^dsRNA^* – ablation; **** p = 3.22E-30; velocity [mm/sec]: *GFP^dsRNA^* – *kst^dsRNA^*: **** p =3.81E-11; *GFP^dsRNA^* – ablation: **** p = 1.54E-05; Peristalsis Efficiency [mm/wave]: *Canton S* – *kst^MIMIC^* / *Df*: **** p = 3.98E-20; *Canton S* – *Trojan ^C^*/ *Df*: **** p = 4.61E-13; *Trojan^C^* – *Trojan^Gal4^* / *Df*: **** p = 1.22E-13; velocity [mm/sec]: *Canton S* – *kst^MIMIC^* / *Df*: **** p = 1.15E-06; *Canton S* – *Trojan^C^/ Df*: **** p = 1.21E-03; *Trojan^C^* – *Trojan^Gal4^* / *Df*: **** p = 5.94E-13.

## Discussion

Here, we present a comprehensive analysis of the ensheathing glia found in the CNS of Drosophila. We developed a split-Gal4 tool to specifically manipulate the ensheathing glia and show for the first time that ensheathing glial cells are highly polarized cells. They function as an internal diffusion barrier with an apical-like cell domain facing towards the neuropil. Polarized membrane lipids as well as several polar localized proteins were identified. The apical sub-membranous cytoskeleton is organized by ß_H_-Spectrin encoded by *karst*. Ensheathing glial cells lacking Karst have an altered cell polarity which is needed for a normal larval locomotor activity. Ensheathing glial cell ablation also leads reduced longevity of adult flies.

To study ensheathing glial cell biology, we utilized the *83E12-Gal4* driver which, as described (Li et al., 2014), appears specific for ensheathing glia. Some of the larval ensheathing glia are diploid and block of cell proliferation reduces their number by 38 %, suggesting that proliferation of these cells contributes to the increase of ensheathing glial complexity. These finding support the notion that some larval ensheathing glia can dedifferentiate, proliferate and redifferentiate to form the more complex adult ensheathing glia (Kato et al., 2020). Ablation of ensheathing glia results in an increased number of adult astrocyte-like cells which suggests that the close interdependence of these two cell types identified in embryonic stages (Peco et al., 2016; Ren et al., 2018) persists to adulthood. Similarly, a plastic interaction also occurs between cortex and ensheathing glia (Coutinho-Budd et al., 2017). Such interactions might also contribute to the functional differences that that appear likely to be associated with for example dorsal or ventral ensheathing glial cells. Unfortunately, no specific molecular tools exist that allow to test this hypothesis. A general role of all ensheathing glial cells is to establish a diffusion barrier around the neuropil, as it was demonstrated by our dye injection experiments. The separation of the neuropil from the cortex in the larva might facilitate the formation of signaling compartments specified by astrocyte-like cells. The 10-fold increase in the number of ensheathing glial cells in the adult brain is likely to further organize neuronal functions into discrete domains (Hadeln et al., 2018; Morris et al., 2007; Omoto et al., 2015; 2018).

As also observed in vertebrates, the formation of signaling compartments can be detected on a gross anatomical level as well as on a circuit level. In both cases, the neuropil-associated glia comprising ensheathing glia are at center stage (Ito et al., 1995) and complexity of regional compartmentalization increases in the adult, mirroring the increase in ensheathing glial cell number (Hartenstein et al., 2008; Spindler and Hartenstein, 2010; Zheng et al., 2018). In vertebrates, a classical example for this fundamental principle can be seen at the rhombomeric organization of the vertebrate hindbrain (Kiecker and Lumsden, 2005). Here, glial cells provide essential functions in boundary formation, too (Yoshida and Colman, 2000). Thus, glia may not only structure the nervous system by compartmentalizing synapses but also act on the level of larger functional units.

The establishment and maintenance of cell polarity is a fundamental cell biological feature and is required for the development of different cell types and tissues. For instance, the mammalian nervous system is protected by the blood-brain-barrier established by polar endothelial cells whereas the blood-brain barrier around the invertebrate nervous system is comprised by polarized glial cells (Carlson et al., 2000; Daneman and Prat, 2015; Limmer et al., 2014; Mayer et al., 2009; Schwabe et al., 2005). Cell polarity is induced by evolutionarily conserved mechanisms (Tepass, 2012; Wodarz and Näthke, 2007). Apical polarity regulators (APRs) comprise a diverse set of proteins centered around the two apical polarity protein complexes Bazooka/Par3 and Crumbs (Riga et al., 2020). In addition, basolateral regulators including the Scribble complex (Scribble, Discs large and Lethal (2) giant larvae and the kinases PAR-1 and LKB1 participate in the definition of a polar cell phenotype (Humbert et al., 2003). Some of these genes (*Par1, Scrib, dlg1*) appear to be expressed by adult ensheathing glia (Davie et al., 2018). For *Scrib* but not *Par1* or *dlg1* this can be confirmed protein trap insertion lines. However, RNAi mediated knockdown of these genes did not cause any morphological defects in larval ensheathing glia. Possibly, similar as in follicle cell development, different mechanisms are in place to guarantee the establishment and maintenance of the polar ensheathing glial cell phenotype (Shahab et al., 2015).

A hallmark of polarized cells is the asymmetric distribution of lipids with PIP_2_ being enriched at the apical surface (Krahn, 2020). To manipulate the ensheathing glial cell polarity, we therefore expressed activated PI3K which should result in a decreased PIP_2_ concentration. Indeed, this led to the formation of ensheathing glial cells lacking PIP_2_ which migrate away from the neuropil and lack β_H_-Spectrin expression (Podogalla, unpublished observations). In turn, reduction in β_H_-Spectrin causes a disrupted PIP_2_ localization. This suggests, that a polar spectrin cytoskeleton might orchestrate lipid composition at the apical membrane domain which subsequently affects polarized distribution of transmembrane proteins such as the Na^+^ / K^+^ ATPase. This might cause to a differential distribution of Na^+^ ions which are used by antiporters involved in metabolite transport. One of them is the Excitatory amino acid transporter 2 (EAAT2) (Featherstone, 2011). Antibodies against EAAT2 are available (Peco et al., 2016) but unfortunately do not allow the analysis of the subcellular localization in larvae. The synaptic neuropil is localized at a unique position within the nerve cord very close to the dorsal surface and thus, the hemolymph. Whereas the ventral as well as the lateral parts of the neuropil are flanked by neuronal cell bodies embedded in cortex glia, only few neuronal cell bodies are found at the dorsal surface of the nervous system. More importantly, no cortex glial cells are found dorsally to the neuropil (Coutinho-Budd et al., 2017; Otto et al., 2018). Thus, metabolites can be transported from the blood-brain barrier to the neuropil only by the ensheathing glia. Upon ablation of the ensheathing glia, the blood-brain barrier forming glial cells might provide enough metabolites to ensure an almost normal function of the nervous system. In case of suppression of Nrv2 expression in the ensheathing glia, ion gradient required for directed transport are not established across the ensheathing glial plasma membrane. At the same time the barrier provided by the ensheathing glia is still in place and therefore metabolically isolates the neuropil which results in an increased mobility possibly aimed to direct the larva to new and better food sources. The neuropil itself is highly organized and its dorsal domain is dedicated to synaptic connections of motor neurons (Landgraf et al., 2003). Possibly, the motor neurons are more sensitive to nutrient supply and thus a more direct metabolic support through the ensheathing glia is required.

## Acknowledgements

We are grateful to B. Altenhein, S. Luschnig, E. Peco and D. van Meyel, and K. Yildirim for generously providing us with antibodies and flies. E. Contreras, M. Krahn, R. Stanewsky, S. Luschnig, S. Schirmeier and S. Rumpf for comments on the manuscript. We thank the Fly EM Project Team at HHMI Janelia and Richard D. Fetter for the Drosophila larval CNS ssTEM volume. We are grateful for all the support of all lab members. This work was supported by the Deutsche Forschungsgemeinschaft through funds to C.K. (SFB 1348, B5).

## Author contributions

N.P. and C.K. designed all experiments. N.P. conducted all experiments. H.K. identified polar expression of Karst, performed initial experiments and together with A.C. analyzed the TEM volume, A.C. contributed data and software, S.Re. and S.Ro. generated constructs used in this study. N.P. and C.K. together with input from all authors wrote the paper.

## STAR Methods

### Material

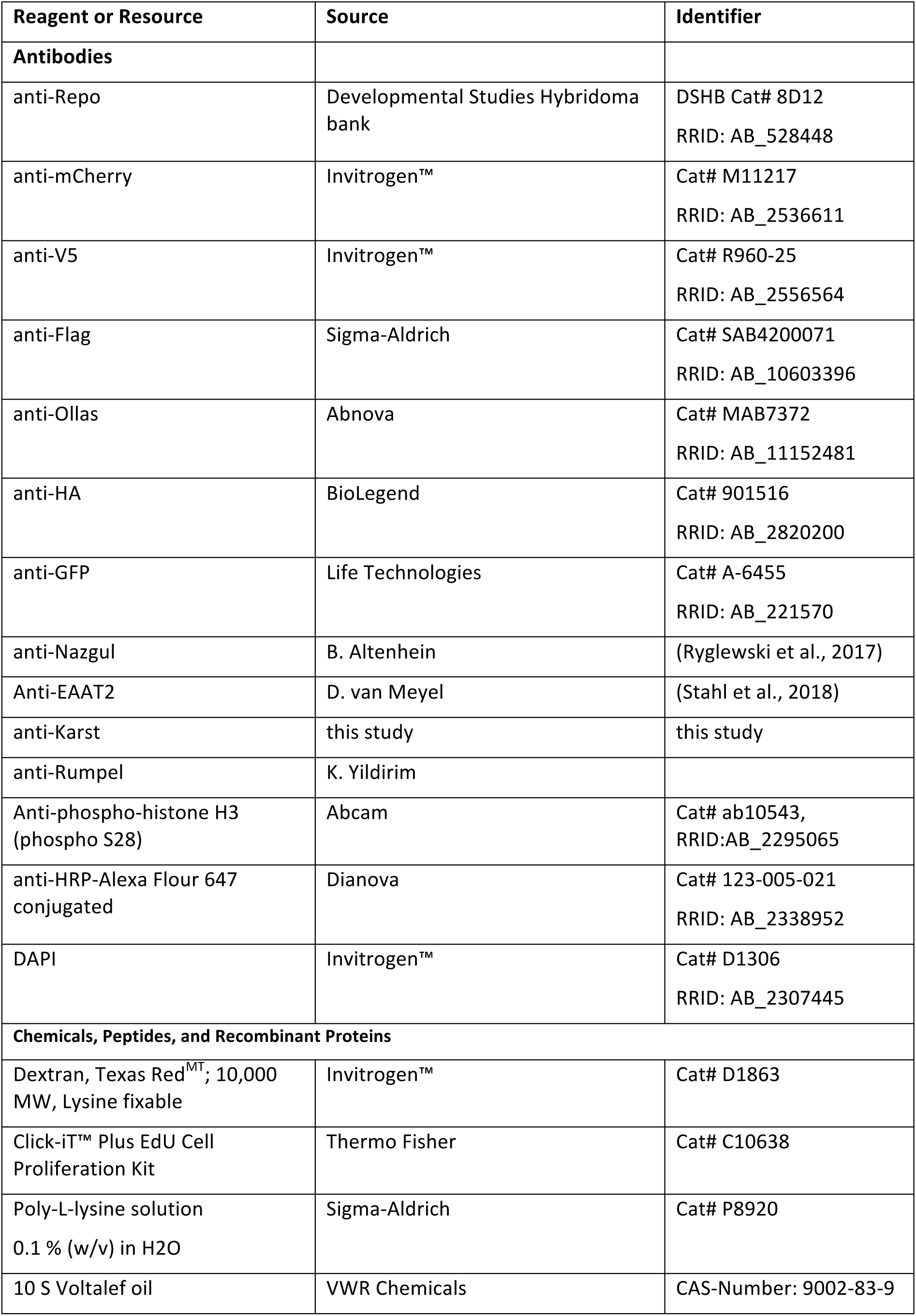
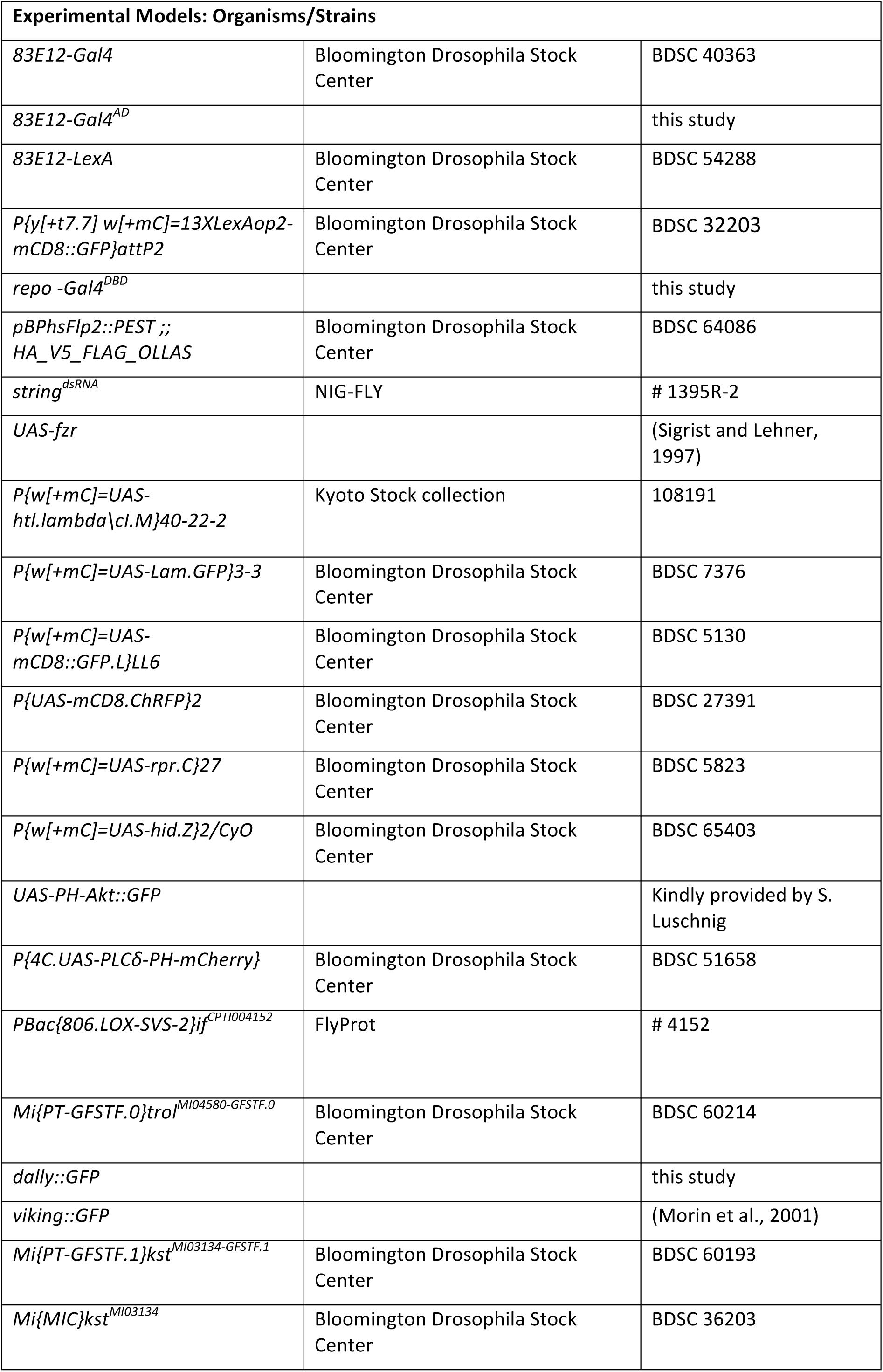
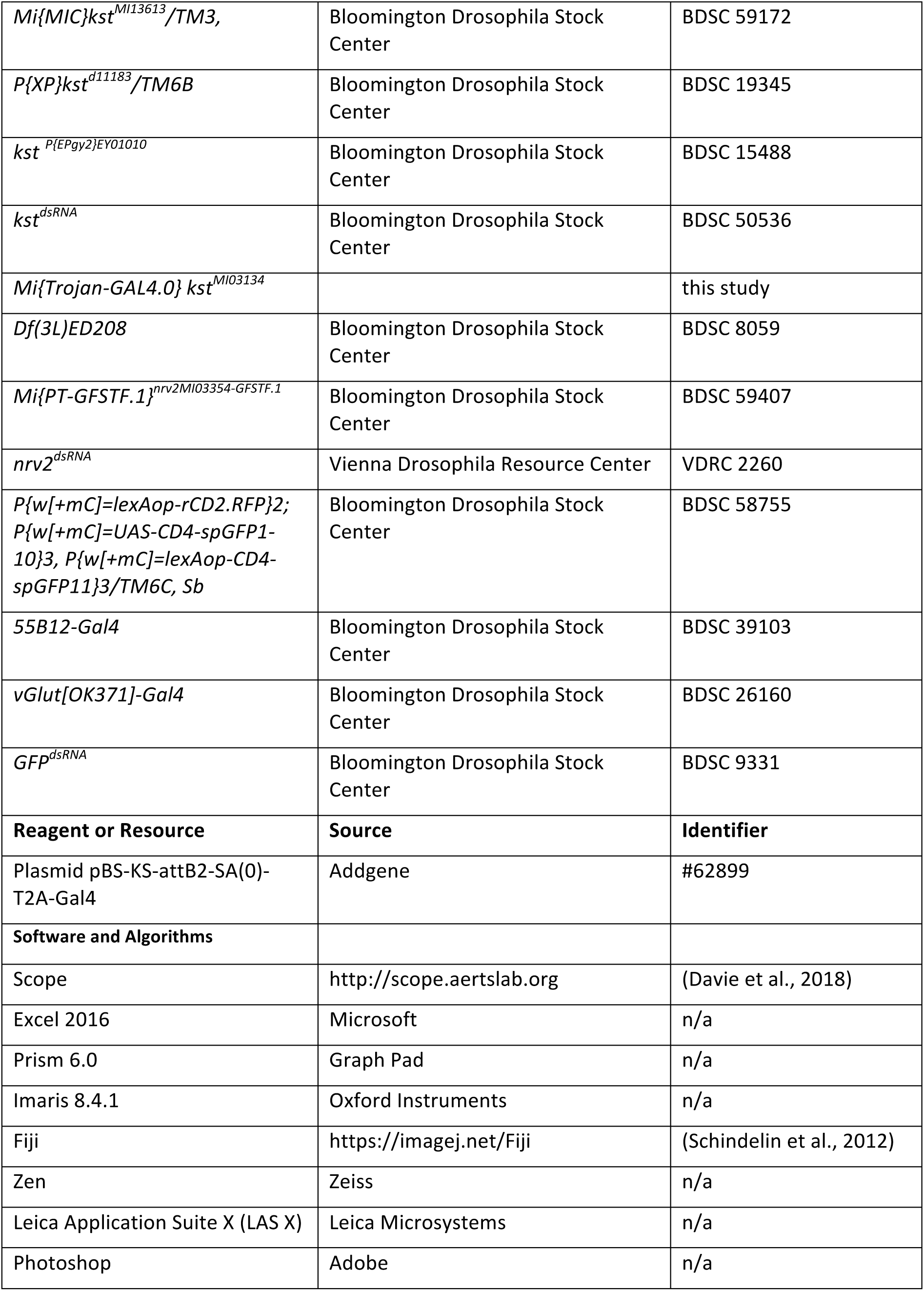

## Methods

### Immunohistochemistry

Immunohistochemistry for larval, pupal and adult brains was performed using standard protocols (Bauke et al., 2015). The following antibodies were used: anti-Repo (8D12; Developmental Studies Hybridoma bank); anti-mCherry (Cat# M11217); anti-V5 (Cat# R960-25) and DAPI (Cat# D1306) (all from Invitrogen™); anti-Flag (Cat# SAB4200071, Sigma-Aldrich); anti-Ollas (Cat# MAB7372, Abnova), anti-HA (Cat# 9015116, BioLegend); anti-GFP (Cat# A-6455, Life Technologies); Anti-phospho-histone H3 (phospho S28) (Cat# ab10543, Abcam); anti-HRP-Alexa Flour 647 conjugated (Cat# 123-005-021, Dianova); anti-Nazgul was a gift from B. Altenhein, Cologne. Rabbit anti-Rumpel was a gift of K. Yildirim, Münster. Guinea Pig anti-EAAT2 was a gift of D. van Meyel. A C-terminally located peptide (^3622^LADERRRAEKQHEHRQN^3639^) shared by all ß_H_-Spectrin proteins was used to immunize rabbits (Pineda, Berlin). For confocal imaging and analysis samples were imaged using a Zeiss LSM 880 or a Leica SP8 confocal microscope using ZEN, Leica LasX and Imaris 8.4.1 Oxford instruments software.

### Electron microscopic analysis

All ssTEM data were generated at the Howard Hughes Medical Institute Janelia Research Campus and processed as described (MacNamee et al., 2016; Ohyama et al., 2015). The CATMAID interface (Saalfeld et al., 2009; Schneider-Mizell et al., 2016) was used to annotate glial cells associated with the neuropil. Glial cell bodies were identified by characteristic color of the cytoplasm and non-neuronal morphology. A cell body node was positioned at the glial soma and annotated using the following information: hemineuromere, glial subtype, serial number for cells in one hemineuromere. The tools built into CATMAID were used to generate 3D renderings and extract image data. Cells identified in RAW data were highlighted using Adobe Photoshop CS6.

### Drosophila melanogaster

Flies were raised according to standard procedure at 25 °C. The following flies were obtained from public stock centers: *83E12-Gal4* (BDSC#40363), *83E12-LexA* (BDSC #54288), P{y[+t7.7] *w[+mC]=13XLexAop2-mCD8::GFP}attP2* (BDSC#32203), *MCFO-2: pBPhsFlp2::PEST;; HA_V5_FLAG_OLLAS* (BDSC#64086), *string^dsRNA^* (#1395R-2), *P{w[+mC]=UAS-htl.lambda\cI.M}40-22-2* (108191), *P{w[+mC]=UAS-Lam.GFP}3-3* (BDSC#7376), *P{w[+mC]=UAS-mCD8::GFP.L}LL6* (BDSC#5130), P{UAS-mCD8.ChRFP}2 (BDSC#27391), *P{w[+mC]=UAS-rpr.C}27* (BDSC#5823), *P{w[+mC]=UAS-hid.Z}2/CyO* (BDSC#65403), *P{4C.UAS-PLCδ-PH-mCherry}* (BDSC#51658), *PBac{806.LOX-SVS-2}if^CPTI004152^* (#4152), *Mi{PT-GFSTF.0}trol^MI04580-GFSTF.0^* (BDSC#60214), *Mi{PT-GFSTF.1}kst^MI03134-GFSTF.1^* (BDSC#60193), *Mi{MIC}kst^MI03134^* (BDSC#36203), *Mi{MIC}kst^MI13613^* (BDSC#59172), *P{XP}kst^d11183^* (BDSC#19345), *kst^P{EPgy2}EY01010^* (BDSC#15488), *kst^dsRNA^* (BDSC#50536), *Df(3L)ED208* (BDSC#8059), , *Mi{PT-GFSTF.1}nrv2^MI03354-GFSTF.1^* (BDSC#59407), *nrv2^dsRNA^* (VDRC#2260), *P{w[+mC]=lexAop-rCD2.RFP}2*; *P{w[+mC]=UAS-CD4-spGFP1-10}3*, *P{w[+mC]=lexAop-CD4-spGFP11}3* (BDSC#58755), 5*5B12-Gal4* (BDSC#39103), *VGlut[OK371]-Gal4* (BDSC#26160), *GFP^dsRNA^* (BDSC#9331). All other fly stocks were generated or were provided by other labs (see list). The MCFO flies were heat-shocked for 1 h at 37 °C in a water bath at different developmental time points as indicated. The Trojan-Gal4 cassettes was exchanged by standard phiC31-integration protocols (Bischof et al., 2007).

### Measurement of DAPI Intensity

To determine the polyploidy of ensheathing glia, we stained larval or adult brains with DAPI, Repo and Elav in order to identify glial cells and neurons. We compared DAPI stained nuclei identified by *83E12-Gal4>UAS-Lam::GFP* with DAPI staining of directly neighboring neuronal nuclei identified by anti-Elav staining. The amount of DAPI staining of neighboring glial and neuronal nuclei was determined using Fiji (Schindelin et al., 2012). In each case the entire nuclear volume was measured.

### EdU incorporation assay

First instar larvae were placed on freshly prepared standard fly food containing 0.2 mM EdU and were kept on the food at 25°C until the third instar stage. Brains of the desired age were dissected and stained for EdU presence using the Click-iT Plus EdU detection kit of Thermo Fisher (# C10638) according the instructions of the manufacture.

### Dye injection Assay

Brains of wandering L3 larvae were dissected on ice and immediately placed on Poly-L-lysine coated object slide and covered by 10 S Volatef halocarbon oil. An automated injection station (FemtoJet 4L, Eppendorf) was used to inject 2.5 mM 10 kDa Texas-Red conjugated dextran (Cat#D1863, Invitrogen) diluted in H_2_O into the neuropil of one brain lobe in each brain. For injections, we used borosilicate glass capillaries of 1 mm outer diameter and 0.58 mm inner diameter (Harvard apparatus 30-0016 capillaries GC100-10) that were pulled on a P-1000 Sutter instrument micropipette puller to reach a tip diameter of about 10 µm as commonly used for DNA injections.

### Longevity Assay

Flies were raised according to standard procedure at 25 °C. The offspring of the control and the ablation experiment were collected for up to 3 days immediately after hatching and separated between males and females. 20 mated females were put in a vial and a total of 200 animals were observed over a period of time. The living flies were counted and placed on fresh food without anesthesia every two days.

### Larval locomotion analysis

Larvae for behavioral experiments were raised at 25 °C according to standard procedure. Larval locomotion of L3 larva was tracked by the FTIR-based Imaging Method (FIM) based on frustrated total internal reflection (FTIR) at room temperature (Risse et al., 2017; 2013). 10-15 larvae per genotype were placed on the tracking arena and after a 60 sec accommodation phase recording started for 3 minutes.

### Quantification and Statistical Analysis

Fiji was used to investigate polar distribution. Cells were divided in two ROI and the fluorescence intensity ratio between apical and basal was compared. Counting of the nuclei and the astrocytes was performed in a 3D model in Imaris 8.4.1 Oxford Instruments. Fluorescence intensities for the Dextran Assay were measured using Fiji (Schindelin et al., 2012) using the normalized fluorescence intensity and the ratio between neuropil and cortex. Statistical analysis and calculation were obtained by Prism 6.0 and Excel.

## Figures

**Supplementary Figure S1 Ensheathing glial cells cover dorsal neurons**

(A) Frontal view of a ventral nerve cord of a third instar larva. Cortex glial cells are labelled using *55B12-Gal4, UAS-CD8::GFP*, the dashed line shows the position of the orthogonal view shown in (B). (B) Orthogonal view, note the absence of cortex glial cell processes dorsally to the neuropil. (C) Frontal view of a ventral nerve cord of a third instar larva with labelled ensheathing glial cells (*83E12-LexA, lexAop-CD8::GFP*), the dashed line shows the position of the orthogonal view shown in (D). (D) Orthogonal view, ensheathing glial cell processes cover the entire neuropil. The arrowheads point towards dorsal cell processes engulfing dorsal neurons. (E,F) GRASP experiment. Larvae with the genotype [*55B12-Gal4 UAS-CD4::GFP^1-10^; 83E12-LexA LexAop-CD4::GFP^11^*]. Expression of GFP^1-10^ is detected by an antibody (in red). Reconstituted GFP is shown in green. Note, that no GFP is reconstituted dorsally to the neuropil. (G-L) Ventral nerve cord of a third instar larva with the genotype: [*OK371-Gal4 UAS-mCherry, 83E12-Gal4 UAS-GFP*]. The dashed line indicates the position of the orthogonal section shown in (H,J,L). (G-J) The morphology of the ensheathing glial cells is shown by GFP staining (green in G,H; white in I,J). Glutamatergic neurons are shown in red (G,H) or in white (K,L). Scale bars are 50µm.

**Supplementary Figure S2 The effect of activated FGF-receptor on ensheathing glia proliferation during pupal stages**

Images of pupal brains dissected at the indicated number of hours (h) after puparium formation (APF). (A,C,E,G,I) Control animals expressing nuclear GFP in the ensheathing glia [*83E12-Gal4 UAS-Lam::GFP*]. Neuronal cell membranes are shown in magenta (anti-HRP staining), ensheathing glia nuclei are in green (GFP). (B,D,F,H,J) Animals expressing activated FGF-receptor together with nuclear GFP in the ensheathing glia [*UAS-!hlt*, *83E12-Gal4 UAS-Lam::GFP*]. Note the predominant increase of ensheathing glia in the thoracic neuromeres (arrowheads). Scale bars are 50µm.

**Supplementary Figure S3 *83E12-Gal4* positive ensheathing glial cell can divide during development**

(A,B) Maximum projections of two ventral nerve cord stained for neuronal cell nuclei (anti-Elav, red), ensheathing glia nuclei (*83E12-Gal4, UAS-Lam::GFP*, anti-GFP, green) and DAPI. (C) 12 examples taken from the maximum projection to illustrate neighborhood relationships. Note, that that the analysis of DAPI signal was conducted all single focal planes with a given nucleus. (D) The ratio of glial and neuronal DAPI intensity is shown for 65 glia / neuron pairs from three brains. Red shading indicates a DAPI intensity ratio of >1.5. (E) Dissected larval CNS with the genotype [*83E12-Gal4, UAS-Lam::GFP*], stained for GFP (green), DAPI (cyan) and phospho-histone H3 (red). (F-H) Dissected pupal CNS of the age indicated with the genotype [*83E12-Gal4, UAS-Lam::GFP*], stained for GFP (green), DAPI (cyan) and phospho-histone H3 (red). Examples of *83E12-Gal4*, phospho-histone H3 positive nuclei are shown in the indicated boxes. (I,J) EdU staining of larval (I) and pupal (J) brains [*83E12-Gal4; UAS-Lam::GFP*]. EdU is shown in red, Lamin::GFP is shown in green to visualize the nuclei of the ensheathing glia. Examples of *83E12-Gal4*, EdU positive nuclei are shown in the indicated boxes. Scale bars are 50 µm.

**Supplementary Figure S4 DAPI staining of larval ensheathing glia**

(A-C) MCFO labeling of an adult brain stained for the expression of V5 (green), HA (red), and FLAG and OLLAS epitopes (blue). *flp* expression (HS) was induced for one hour, during third instar larval stage (A,B) or at the onset of puparium formation (0h APF). Note that not all ensheathing glia that are present in the adult CNS are labelled when *flp* is expressed in early development. D) Third instar larval brain with ensheathing glia labelled using *83E12-Gal4* driving membrane bound GFP. (E) Similar aged larval brain with ensheathing glia labelled using the split-Gal4 combination driving membrane bound GFP [*83E12-Gal4^AD^*, *repo-Gal4^DBD^*, UAS-CD8::GFP]. (F,G) Adult brains of animals carrying either the *83E12-Gal4* or the split-Gal4 combination. (H-J) Ablation of ensheathing glia. Animal of the genotype [*83E12-Gal4^AD^, repo-Gal4^DBD^, UAS-hid, UAS-CD8::GFP*] lacks detectable GFP-expression in the CNS. Few wrapping glial cells along the peripheral nerve are not ablated (asterisk). The dashed white line indicated the position of the orthogonal section shown in (J). [Scale bars are: larval CNS 50 µm, adult CNS 100 µm.

**Supplementary Figure S5 Polar organization of the adult ensheathing glia**

(A) Coexpression of *PH-AKT-GFP* and *PH-PLCδ-mCherry* in adult ensheathing glia (EG). The boxed areas are shown in higher magnification in (B-E). Note that green fluorescence indicating PIP_3_ is preferentially seen towards cortical regions (arrowheads), whereas magenta staining indicating PIP_2_ is preferentially found facing the neuropil (arrows). Scale bar is 100 µm.

**Supplementary Figure S6 Expression of extracellular matrix components in the adult brain**

The figure shows SCENIC representations of the 57K scRNA seq data set of the Aerts laboratory (Davie et al., 2018). SCope analysis for the genes indicated in each top right corner is shown. Each dot represents a single cell. The color coding indicates the expression level. Red: strong expression, black: low expression, grey: no expression. Expression of the transcription factor Repo defines the glial complement, expression of further markers allows the definition of glial subtypes (Davie et al., 2018). *inflated* (*if*) is expressed in ensheathing glia, astrocyte-like glia and perineurial glia. The genes encoding the ECM components *trol* (Perlecan), *vkg* and *col4a1* (CollagenIV), *dally* (heparansulfate proteoglycan), *Tigrin* and *SPARC* are strongly expressed in ensheathing glia (arrowheads). The genes encoding the different Laminin subunits (*LanA*, *LanB1*, *LanB2*) are weakly expressed by glia.

**Supplementary Figure S7 Karst expression**

(A-D) Silencing of Karst^GFP^ expression in larval brains. (A) Single focal plane of a CNS with the genotype [*repo-Gal4, UAS-GFP^dsRNA^, kst^MiMIC::GFP^*]. The position of trachea is indicated (asterisks). The dashed line shows the position of the orthogonal section shown in (B). The arrows point to unspecific binding of the anti-GFP antibody to the outer surface of the CNS. (C,D) Single focal plane of a CNS with the genotype [*moody-Gal4, UAS-GFP^dsRNA^, kst^MiMIC::GFP^*]. Note, that GFP expression is still found around the neuropil (arrowheads). (E) Stage 16 control embryo and (F) *kst* deficient embryo stained for Karst localization. To obtain specific antibodies, rabbits were immunized using a short peptide (^3622^LADERRRAEKQHEHRQN^3639^) shared by all ß_H_-Spectrin proteins. The purified antiserum was used to stain control and *karst* null mutant embryos as indicated. Scale bars are: 50 µm.

**Supplementary Figure S8 Karst expression in ensheathing glia declines during pupal development**

Differentially aged pupal brains with the hours after puparium formation (APF) indicated were stained for Karst::GFP (green), CD8::mCherry expression driven by *83E12-Gal4* (magenta) and HRP (blue) to label all neuronal membranes. The positions of the orthogonal sections are indicated by dashed lines. (A-F) *Karst::GFP* expression can be detected up to 72h APF. (G,H) In 96 h APF old pupae, ensheathing glia reorganize and Karst expression disappears. Scale bar is 100 µm.

**Supplementary Figure S9 Opposite effects of *karst* and *nrv2* on larval locomotion**

(A-C) Representative larval locomotion tracts of the genotypes indicated. (D) Quantification of peristalsis efficiency. Note that knockdown of *nrv2* specifically in ensheathing glia results in an increase in the peristalsis efficiency [*83E12-Gal4^AD^, repo-Gal4^DBD^, nrv2^dsRNA^*], whereas knockdown of *karst* results in a decrease [*83E12-Gal4^AD^, repo-Gal4^DBD^, kst^dsRNA^*]. (G) Quantification of crawling velocity. Knockdown *nrv2* leads to an increase in the crawling velocity, whereas knockdown of *karst* results in a decrease. (H) Quantification of bending distribution. Quantification Wilcoxon rank-sum test, n=50. Peristalsis Efficiency [mm/wave]: *GFP^dsRNA^* – *kst^dsRNA^*: **** p = 3.22E-30; *GFP^dsRNA^ – nrv^dsRNA^*: **** p = 1.20E-17; velocity [mm/sec]: *GFP^dsRNA^* -*kst^dsRNA^*: **** p = 3.81E-11;; *GFP^dsRNA^ – nrv^dsRNA^*: **** p = 8.49E-05.

**Supplementary movie 1**

Dye penetration in control third instar larvae

**Supplementary movie 2**

Dye penetration in ensheathing glia ablated third instar larvae

