## Supplementary Figures for "Drosophila ß_Heavy_-Spectrin is required in polarized ensheathing glia that form a diffusion-barrier around the neuropil"

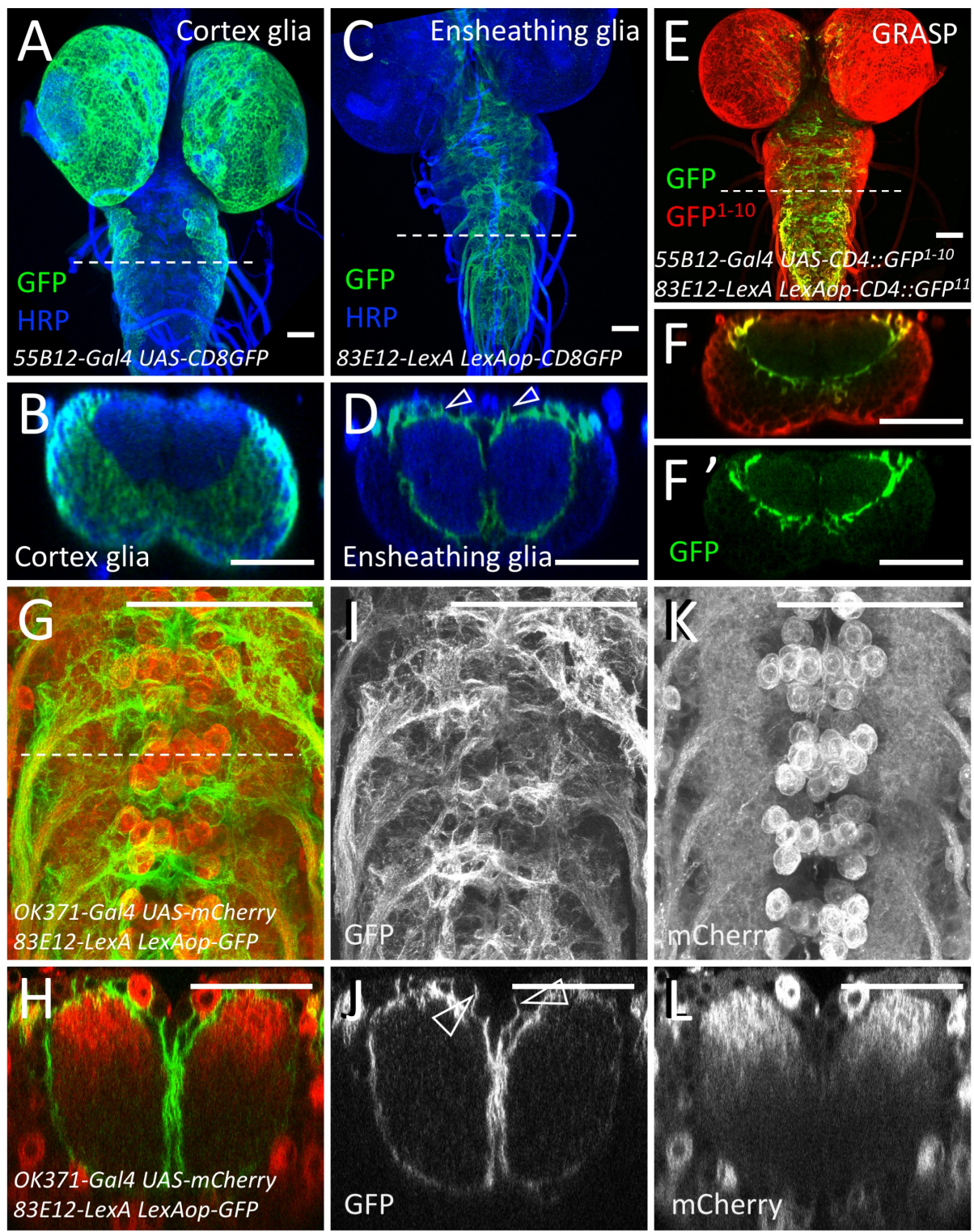

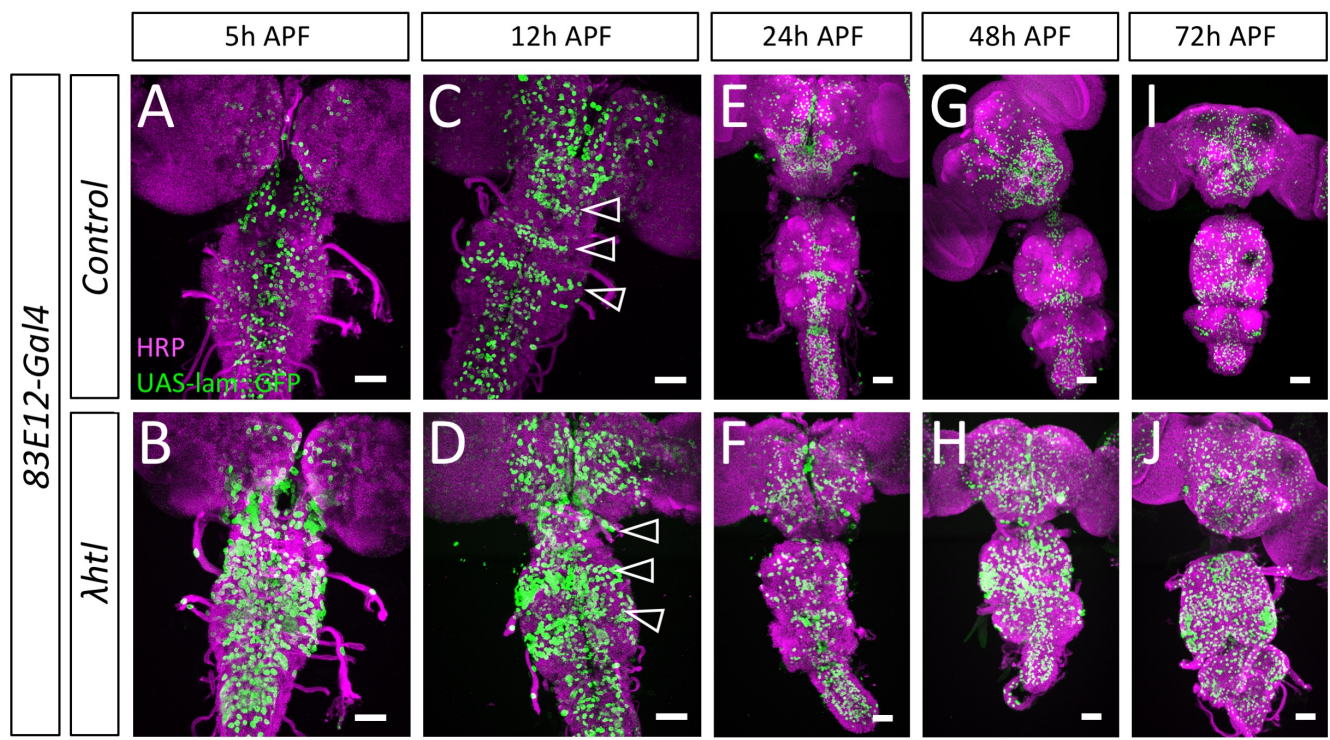

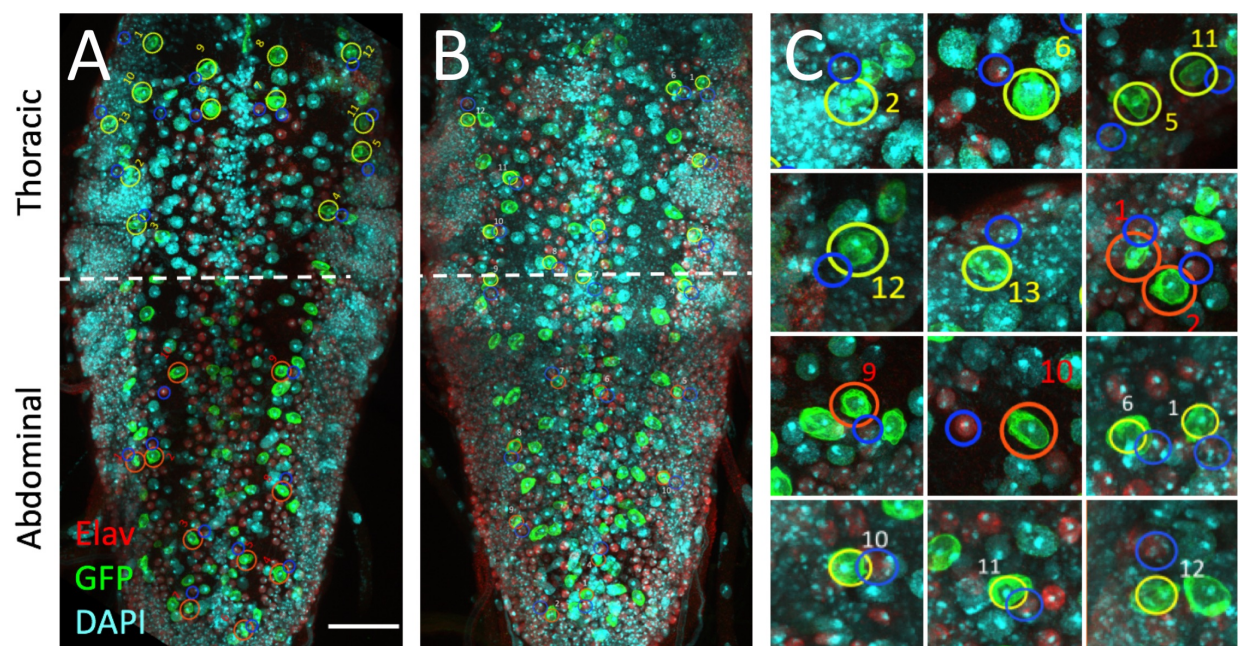

### D DAPI ratio

| Thoracic | Nucleus pair # | 1 | 2 | 3 | 4 | 5 | 6 | 7 | 8 | 9 | 10 | 11 | 12 | 13 |
| --- | --- | --- | --- | --- | --- | --- | --- | --- | --- | --- | --- | --- | --- | --- |
| Glia/Neuron | Brain 1 | 1,3 | 1 | 1 | 0,8 | 1 | 1,9 | 1,8 | 1,5 | 1,8 | 1,5 | 1,5 | 1,3 | 1,4 |
|  | Brain 2 | 1,4 | 1,4 | 1,3 | 1,5 | 1,2 | 1,9 | 1,4 | 1,4 | 1,1 | 1,6 | 1,6 | 1,2 |  |
|  | Brain 3 | 1,7 | 2,5 | 1,5 | 1,7 | 1 | 1 | 1,4 | 0,9 | 0,9 | 2,1 |  |  |  |

| Abdominal | Nucleus pair # | 1 | 2 | 3 | 4 | 5 | 6 | 7 | 8 | 9 | 10 |
| --- | --- | --- | --- | --- | --- | --- | --- | --- | --- | --- | --- |
| Glia/Neuron | Brain 1 | 2 | 1,8 | 1,9 | 2,3 | 1,7 | 2,2 | 2,4 | 2 | 2 | 1,9 |
|  | Brain 2 | 2 | 2,3 | 1,7 | 2,2 | 1,9 | 1,9 | 2 | 1,7 | 2,3 | 2,1 |
|  | Brain 3 | 1,9 | 2,1 | 2 | 2,4 | 3,2 | 1,8 | 1,9 | 1,5 | 2 | 2,1 |

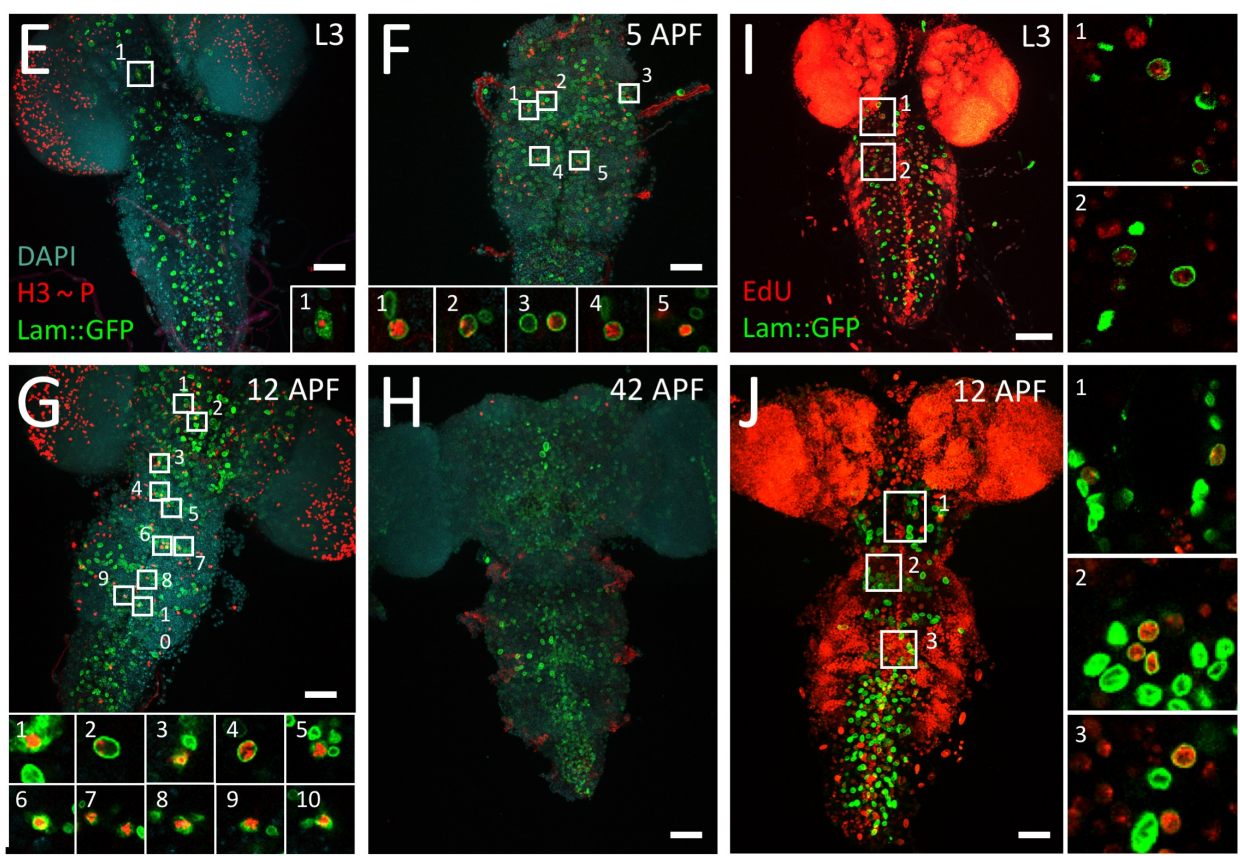

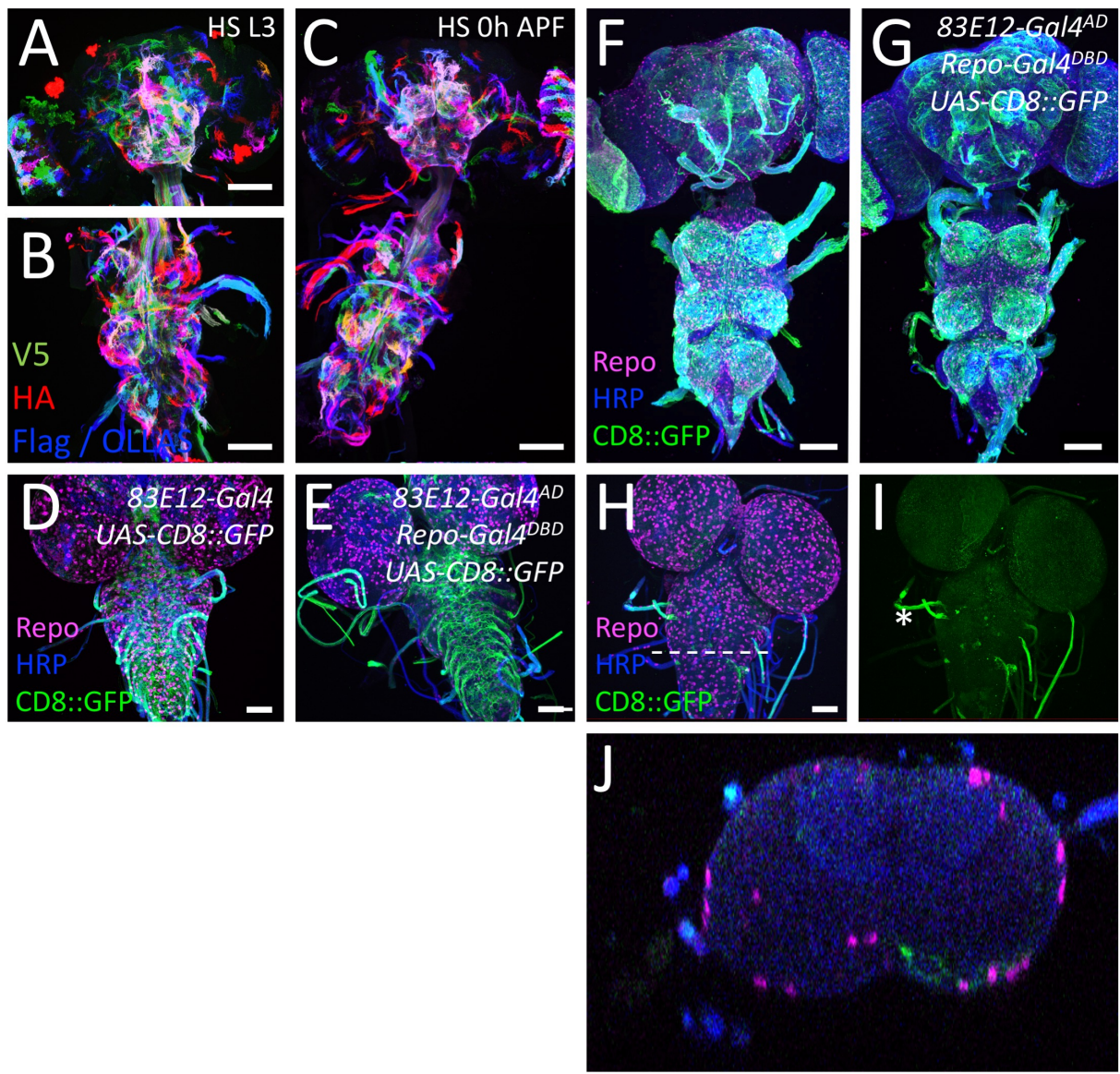

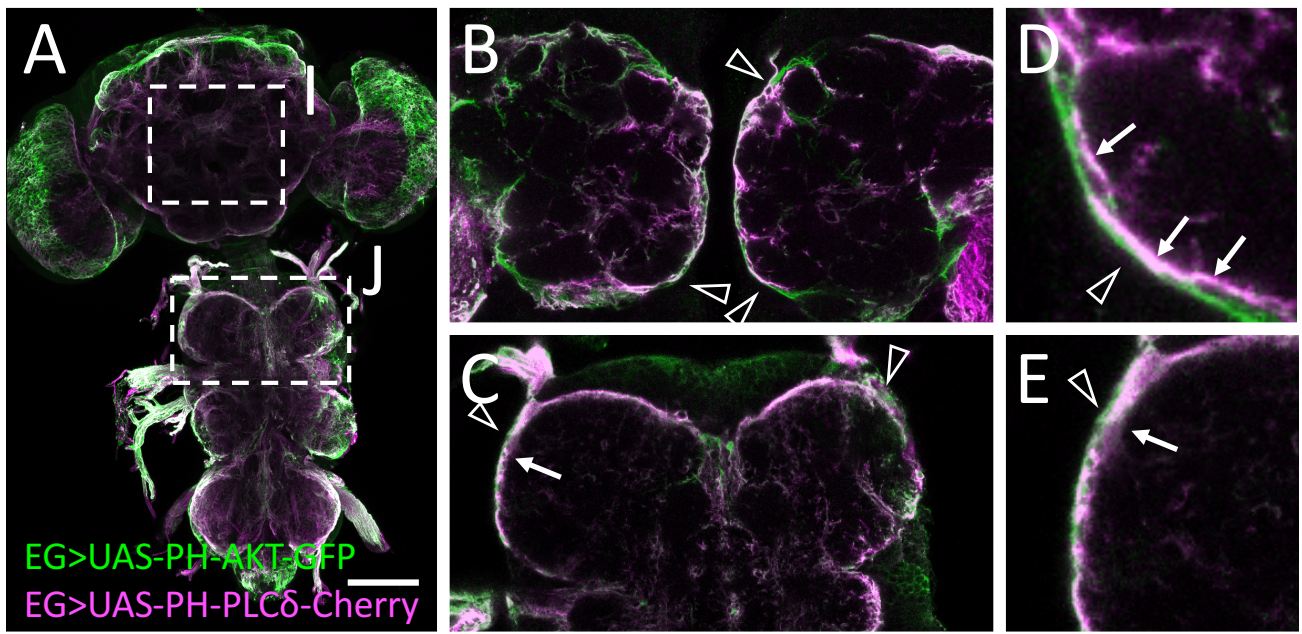

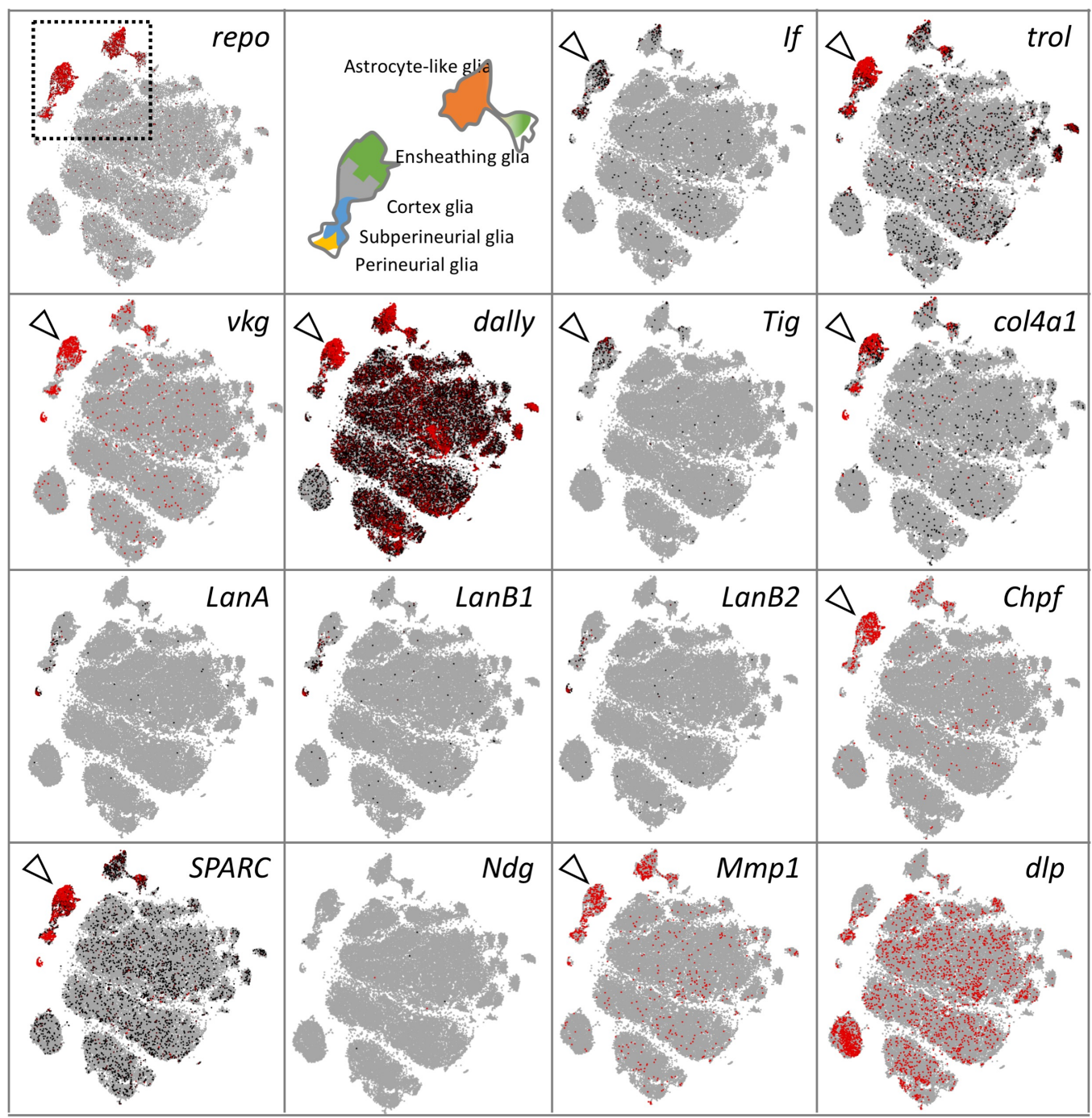

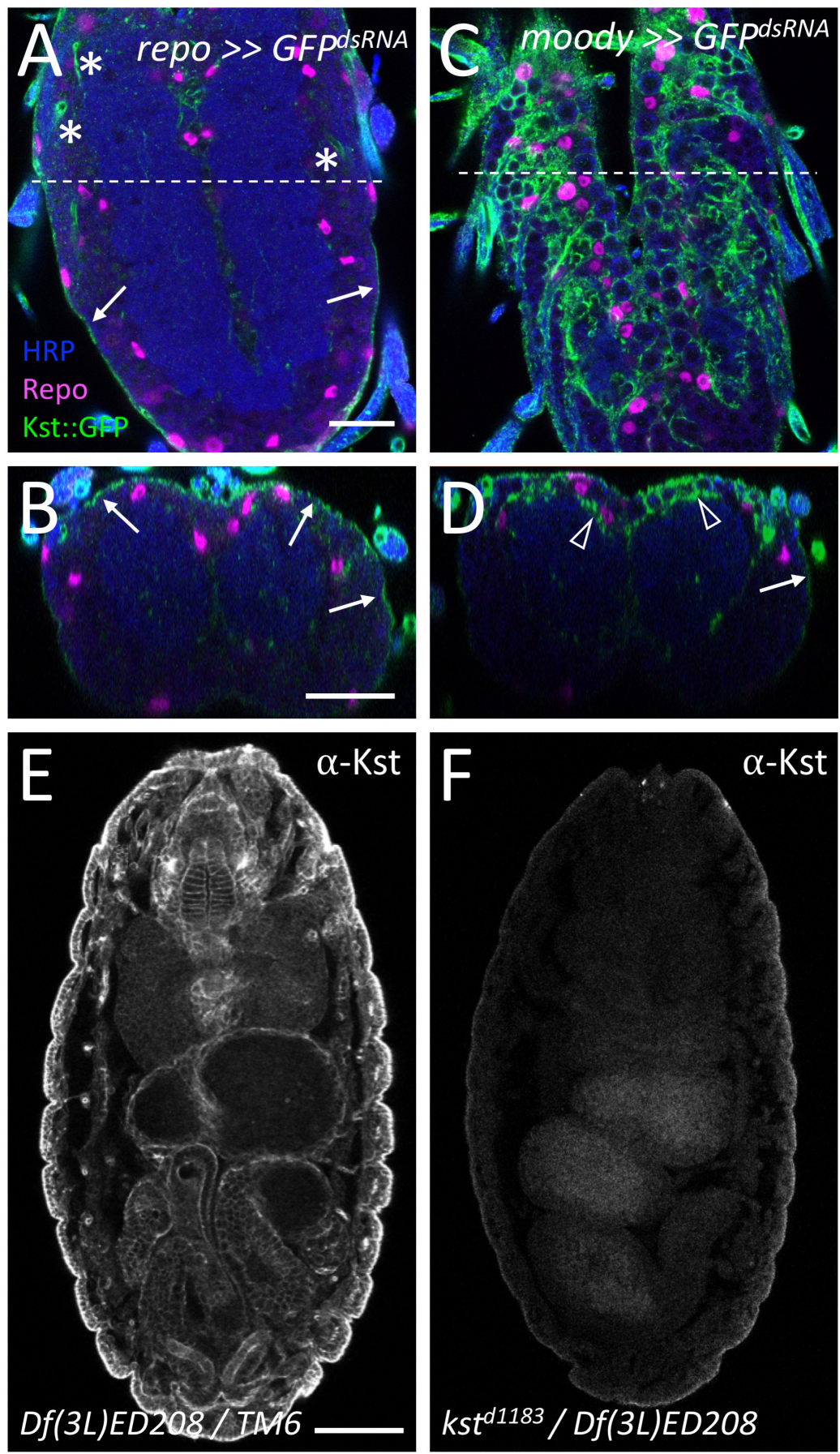

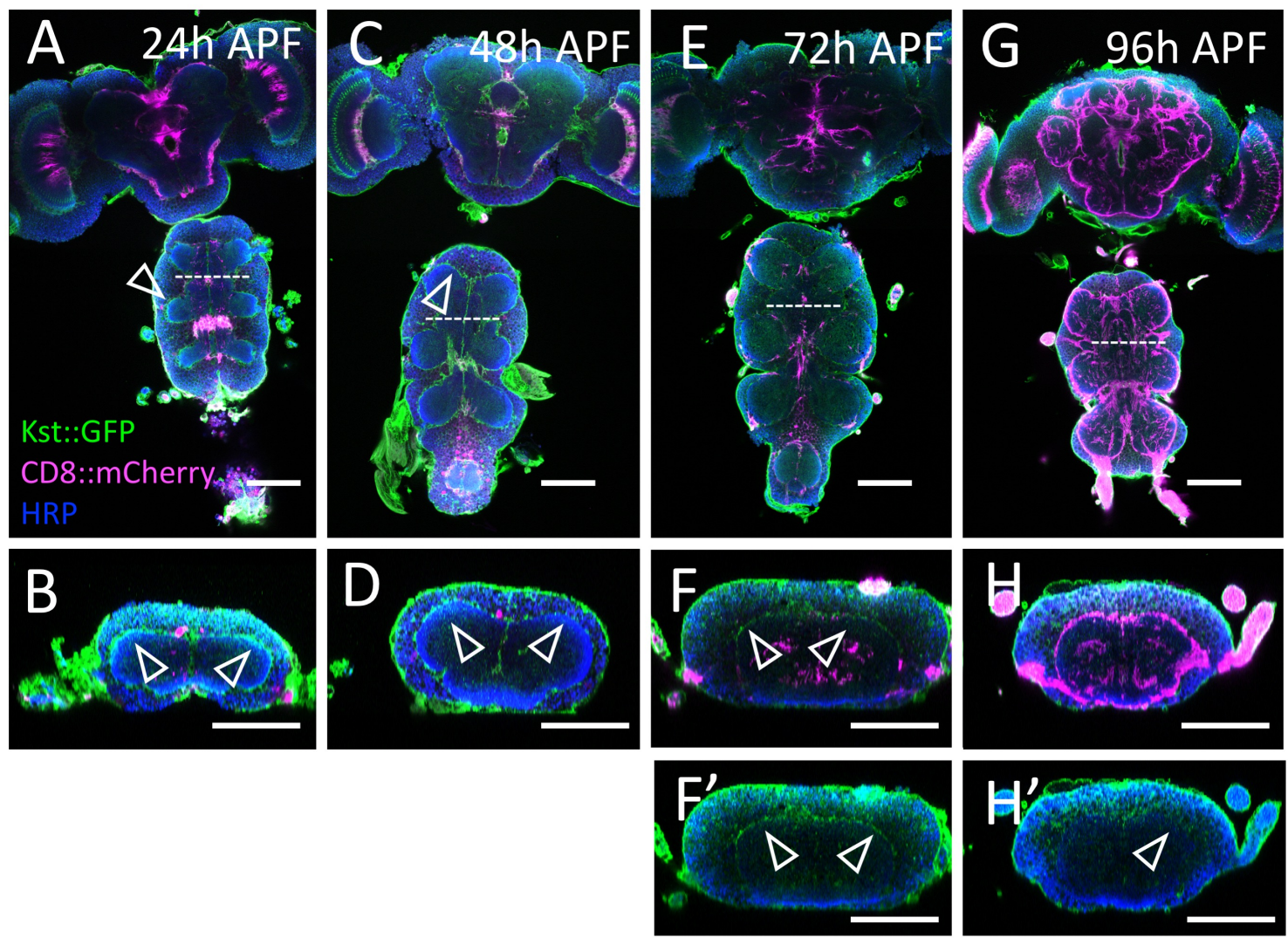

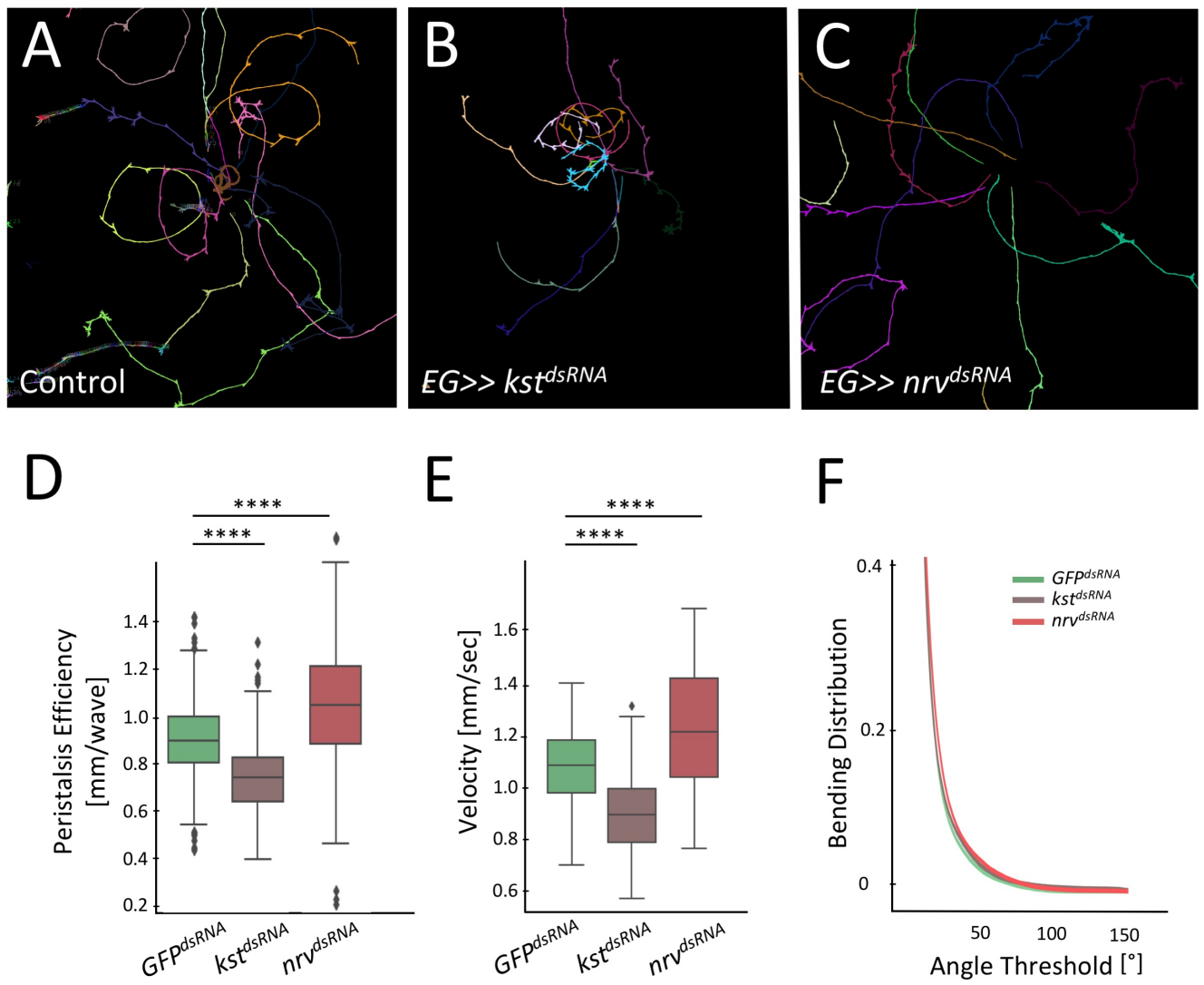
